# Distinct Inhibitory Pathways Control Velocity and Directional Tuning in the Retina

**DOI:** 10.1101/2022.01.13.476257

**Authors:** Mathew T. Summers, Marla B. Feller

## Abstract

The sensory periphery is responsible for detecting ethologically relevant features of the external world, using compact, predominantly feedforward circuits. Visual motion is a particularly prevalent sensory feature, the presence of which can be a signal to enact diverse behaviors ranging from gaze stabilization reflexes, to predator avoidance or prey capture. To understand how the retina constructs the distinct neural representations required for these diverse behaviors, we investigated two circuits responsible for encoding different aspects of image motion: ON and ON-OFF direction selective ganglion cells (DSGCs). Using a combination of 2-photon targeted whole cell electrophysiology, pharmacology, and conditional knockout mice, we show that distinct inhibitory pathways independently control tuning for motion velocity and motion direction in these two cell types. We further employ dynamic clamp and numerical modeling techniques to show that asymmetric inhibition provides a velocity-invariant mechanism of directional tuning, despite the strong velocity dependence of classical models of direction selectivity. We therefore demonstrate that invariant representations of motion features by inhibitory interneurons act as computational building blocks to construct distinct, behaviorally relevant signals at the earliest stages of the visual system.

## Introduction

Rather than acting as a simple camera forming a pixel-by-pixel map of image luminance, the mammalian retina is comprised of diverse arrays of feature detectors, each encoding distinct components of the visual scene such as color, oriented edges, or motion^1–3^. These output channels convey visual information to differing brain regions to mediate appropriate behaviors, such as pupillary light reflexes, looming responses, or optokinetic reflexes^4^. Directional image motion is a particularly ubiquitous feature which is encoded by an estimated 35% of the mouse retina’s output neurons^5^. However, different DSGC types meet different behavioral demands and thus encode different types of motion. ON DSGCs project to the accessory optic system, and mediate gaze stabilizing reflexes by encoding low velocity, global motion^6–9^. ON-OFF DSGCs target the image-forming brain regions of the superior colliculus and lateral geniculate nucleus, and encode local object motion across a broad range of velocities^10–12^. Thus the retina computes at least two parallel output channels for motion direction, each of which contains different information on the velocity of image motion. How the retina constructs distinct representations of sensory features using a limited pool of largely feedforward interneurons remains an active area of research^13–17^.

Tuning for direction and velocity are linked by circuit models for elementary computations of direction. Early theory work to identify the minimum computations necessary for directional motion detection conceptualized circuit models that compute spatiotemporal correlations by comparing spatially offset luminance signals with some time delay or lowpass filter. The Hassenstein-Reichardt correlator and Barlow-Levick rectifier are instantiations of these correlation computing models, and though they differ in implementation, both rely on some form of temporal offsetting to compare signals^18–20^ (Fig. 1a). Consequently, directional responses are only expected when the displacement of a stimulus aligns with the temporal offsetting of the model, establishing a tacit link between direction and velocity tuning. Many direction selective neurons show this tuning dependence upon velocity, including J-RGCs, F-mini RGCs, and the GABA-independent computation of Hb9 DSGCs, as well as cortical visual neurons^2,21–23^. ON DSGCs are velocity tuned^6,13^ and may therefore follow these circuit models. In contrast, ON-OFF DSGCs are directionally tuned largely independent of velocity^11^. Whether this velocity invariance is due to a combination of mechanisms implemented at different velocities remains to be determined.

**Figure 1.**
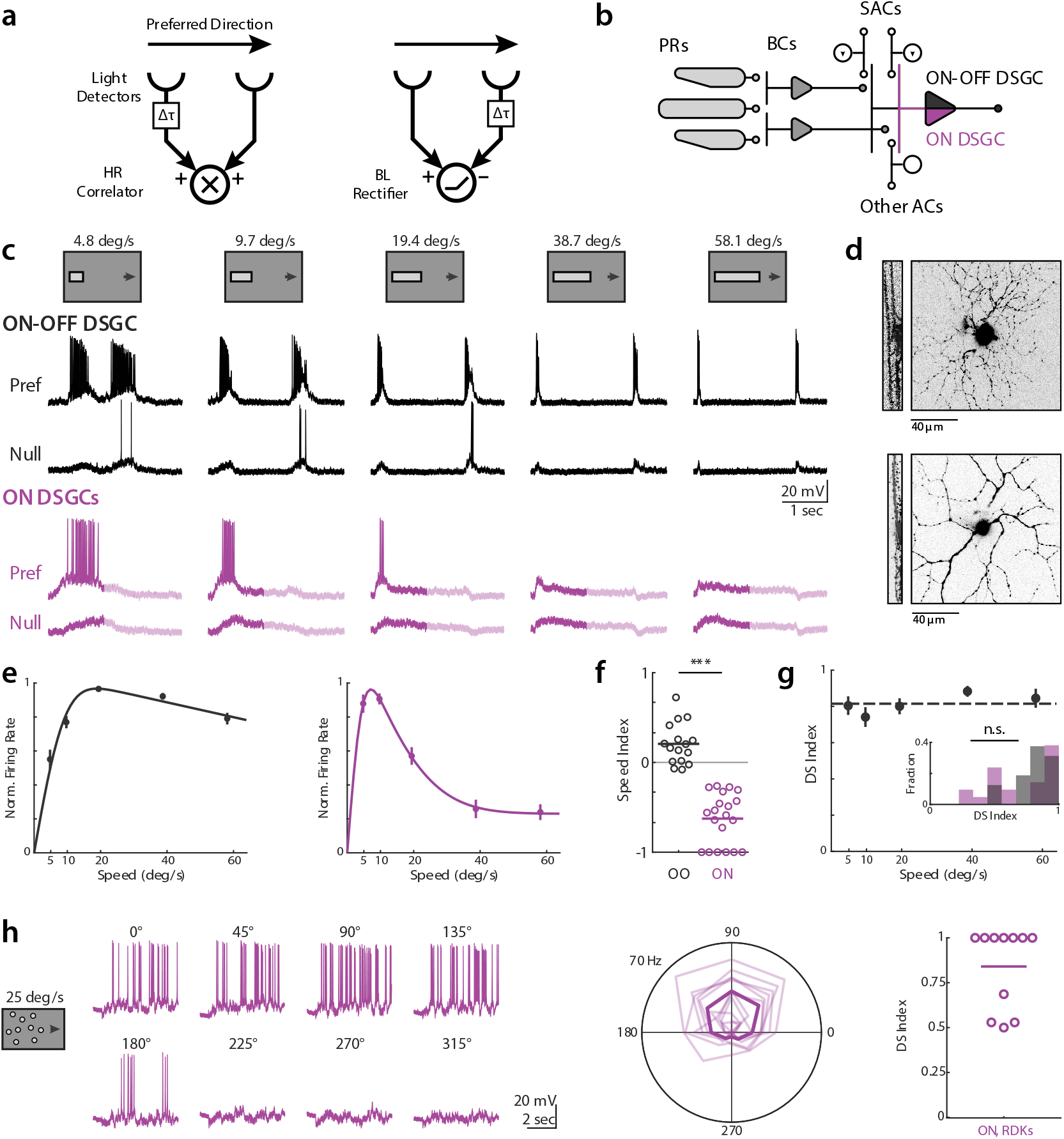
ON-OFF and ON DSGCs are directionally tuned independent of velocity. (a) Mechanistic models of direction selectivity, (*left*) Hassenstein-Reichardt Correlator and (*right*) Barlow-Levick Rectifier. Both models depend upon comparing signals from spatially offset subunits with a differential time delay Δt. (b) Schematic of ON-OFF exclusive (*black*) and ON shared (*purple*) direction selective ganglion cells (DSGCs) circuit components. PRs; photoreceptors. BCs; bipolar cells. ACs; amacrine cells. SACs; starburst amacrine cells. (c) Example ON-OFF (*black*) and ON (*purple*) DSGC current clamp recordings for elongated bar stimuli where bar length scales with speed. Opaque lines shows analysis window restricted to On responses. (d) Example ON-OFF (*top*) and ON DSGC (*bottom*) TexasRed cell fills, showing xy projection (*right*) and xz ON and OFF layer bistratification (*left*). (e) Population-averaged velocity tuning curves of normalized peak firing rate. Error bars show standard error of the mean, solid lines show fits to an integral over difference of Gaussians. (f) Speed indices of current clamp spiking data for ON-OFF (*black*) and ON DSGCs (*purple*), compared via two-sided Wilcoxon rank-sum test; \*\*\**P* = 3.6 × 10^−7^, 16 ON-OFF DSGCs in 11 mice, 21 ON DSGCs in 17 mice. (g) Population-averaged velocity tuning curve of DSI in ON-OFF DSGCs. Dashed line shows velocity-averaged DSI. Inset shows DSI histogram comparing velocity-averaged ON-OFF DSGC DSIs with ON DSGC DSIs at the velocities for which their firing rate is highest. Comparison was made via two-sided Wilcoxon rank-sum test, NS, *P* = 0.60, 16 ON-OFF DSGCs in 11 mice, 21 ON DSGCs in 17 mice. (h) (*left*) Example ON DSGC current clamp recordings for random dot kinetogram (RDK) stimuli moving coherently in one of eight directions. (*Middle*)Polar plot directional tuning curves of ON DSGCs spiking for RDK stimuli. Preferred directions are aligned to 90 degrees. Transparent lines are tuning of individual cells, bold line is population average. (*Right*) Direction selective indices computed from peak firing rate.

Recently, several circuit mechanisms have been identified to contribute to DSGC encoding of motion direction. A critical source of directional tuning is inhibitory input from starburst amacrine cells (SACs), which provide greater inhibition for null relative to preferred direction motion^24^. This directional inhibition is the product of two asymmetries: individual SAC processes are themselves tuned for motion outward from the SAC soma, and SAC processes preferentially synapse onto DSGCs with preferred directions that are antiparallel to the SAC dendrite^25^. Preference for outward motion in the SAC process itself appears to independent of velocity^26^, and may therefore be a source of velocity invariant tuning to both ON and ON-OFF DSGCs. However, other mechanisms have also been described to contribute to direction selectivity, including directional excitation seen in both ON and ON-OFF DSGCs^27–29^, and spatially offset inhibition^30,31^, which has been reported to facilitate ON-OFF DSGC tuning by introducing differential timing offsets between excitation and inhibition for preferred versus null motion. Independent, local computations within DSGC dendrites due to the subcellular arrangement of excitatory and inhibitory synapses could also contribute to directional tuning^32,33^. The relative contributions of these various mechanisms, and whether some show the strong velocity dependence predicted by classical circuit models, has not been tested.

Here we use a combination of voltage clamp measurements, dynamic clamp, and conductance modeling to show that directional inhibition is the dominant source of direction selectivity across physiological velocities, and that this tuning is imparted in a velocity invariant manner to both ON and ON-OFF DSGCs. We further use conditional knockout mice to show that velocity invariant inhibition is inherited entirely from SACs, and pharmacology to confirm that ON DSGC velocity preference is the product of non-directional glycinergic inhibition. These findings support a model of distinct inhibitory circuits mediating tuning for direction and velocity among retinal feature detectors.

## Results

### ON and ON-OFF DSGCs are robustly tuned for direction across velocities

To study the relative directional computations of ON and ON-OFF DSGCs, we performed 2-photon targeted current clamp recordings using Hoxd10-GFP mice to label ON DSGCs, and a combination of Hoxd10-GFP, Drd4-GFP, and Trhr-GFP mice to label ON-OFF DSGCs^12,34,35^. We set out to perform a systematic analysis of the directional tuning of ON and ON-OFF DSGCs across a range of speeds that spanned the lower bound of optokinetic reflex tuning to the upper end of saccade-like velocities^36,37^ (Fig. 1b, d). To rigorously compare the tuning of ON and ON-OFF DSGCs, we used elongated drifting bar stimuli to isolate the initial ON response of each cell type (Fig. 1c). We scaled the length of bar stimuli with velocity to ensure separation of ON and OFF responses, and accordingly restricted our analysis to the ON response. Consistent with previous reports^6,13^, we found ON DSGCs responded strongly at low velocities (~5-10 °/sec) but weakly if at all at higher velocities, while ON-OFF DSGCs were broadly responsive over a range of physiological speeds, with slight preference for moderate to high velocities (~20-60 °/sec) (Fig. 1e). Spatially restricted drifting gratings recapitulated the tuning of our elongated bar stimuli, ruling out potential artifacts due to greater surround suppression for our high speed bars (Extended Fig. 1).

We used tuning indices to quantitatively compare the selectivity of ON and ON-OFF DSGCs. To quantify velocity preference, we used a speed index ranging from −1 to +1 to denote respective tuning for low vs high speeds, and with 0 indicating no preference between speeds. As expected, the speed tuning preferences of ON and ON-OFF DSGCs were significantly different by this metric (Fig. 1f, Table 1). Directional tuning was quantified with a direction selectivity index (DS index) ranging from +1 to 0, respectively indicating complete or no preference for preferred over null direction motion. ON DSGC direction selectivity was difficult to assess at high velocities due to minimal spiking activity, but ON-OFF DSGC directional selectivity was largely speed invariant (Fig. 1g). We found no significant difference when comparing DS indices of ON DSGCs at velocities eliciting peak firing with the DS indices of ON-OFF DSGCs averaged across the range of speeds tested, indicating both cell types are comparably tuned when active.

Previous work showed that random dot kinetograms (RDKs) elicit low gain optokinetic reflexes in mice, even when the component dots move at relatively high speeds^38^. Given the low velocity preference of ON DSGCs which are thought to mediate optokinetic reflexes, we sought to investigate the responsiveness of ON DSGCs to RDK motion. We found that ON DSGCs responded robustly and in a directionally tuned manner to RDK motion at 25 °/sec, indicating the operation of a directional tuning mechanism at velocities above which ON DSGCs are normally responsive (Fig. 1h).

### Synaptic inputs underlying tuning for direction and velocity

To investigate the synaptic origins of DSGC spike tuning we performed voltage clamp recordings in each cell type while using our elongated bar stimuli. ON-OFF DSGCs received phasic inhibitory postsynaptic currents (IPSCs) both at the entrance and exit of the bar into the DSGC’s receptive field; these IPSCs were directionally tuned, and largely unchanged at different velocities (Fig. 2a). ON DSGCs received large, phasic IPSCs at bar onset, and frequently had smaller magnitude currents at bar offset, perhaps consistent with a previous report of ON DSGC dendrites partially arborizing in OFF sublaminae^34^. We restricted our analysis to ON responses in both cell types to allow for direct comparisons across cell types. While ON DSGC IPSCs also showed directional tuning, the magnitude of IPSCs increased dramatically with velocity, unlike ON-OFF DSGCs (Fig. 2a).

**Figure 2.**
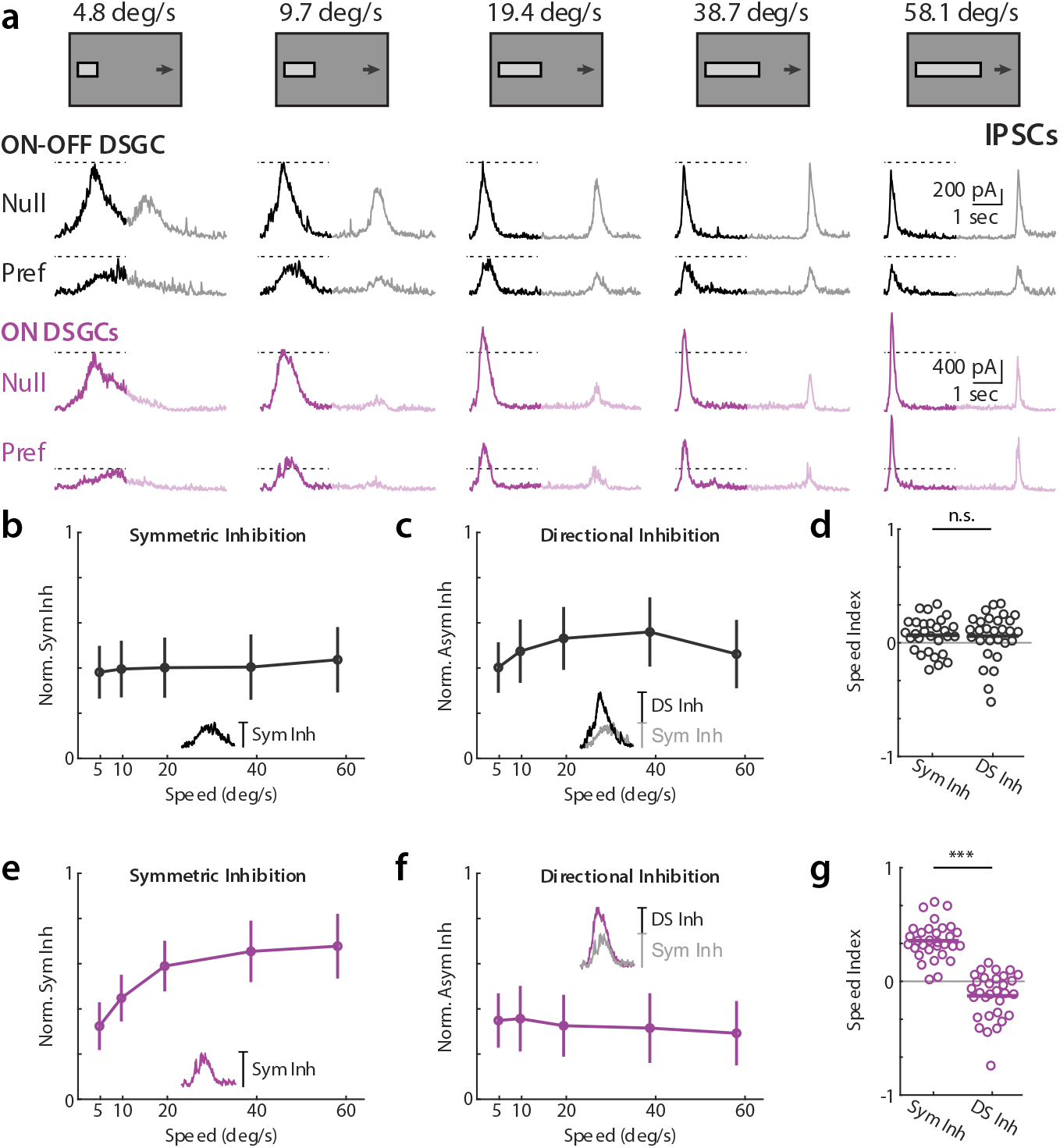
ON-OFF and ON DSGCs receive velocity invariant directionally tuned inhibition and ON-DSGCs receive symmetric velocity tuned inhibition. (a) Example ON-OFF (*black*) and ON (*purple*) DSGC IPSC recordings for drifting bar stimuli. Opaque traces shows analysis window restricted to ON responses. Dashed lines indicate IPSC amplitude at the lowest tested velocity. (b) Population-averaged velocity tuning curves of symmetric inhibition normalized to each cell’s maximal null direction IPSC. Error bars show standard deviation. Inset shows measurement of symmetric inhibition as amplitude of preferred direction IPSC. (c) Population-averaged velocity tuning curves of directional inhibition normalized to each cell’s maximal null direction IPSC. Error bars show standard deviation. Inset shows measurement of directional inhibition as amplitude difference of null minus preferred direction IPSC. (d) Speed indices of symmetric and asymmetric inhibition. IPSC speed indices were compared via two-sided Wilcoxon signed-rank test; NS, *P* = 0.92, 29 cells in 19 mice. (e-g) Same as b-d, but for ON DSGCs. ON DSGC IPSC speed indices were compared via two-sided Wilcoxon signed-rank test; \*\*\**P* = 1.6 × 10^−6^, 31 cells in 21 mice.

ON and ON-OFF DSGCs are known to receive synaptic input from SACs, as well as non-SAC amacrine cells^9,31,39–41^. In order to assess the specific contribution of inhibitory inputs to directional tuning at each velocity, we interpreted IPSCs as being comprised of symmetric and directionally tuned components. We reasoned that the magnitude of inhibition elicited by preferred direction motion constituted an upper bound for non-directional input, and thus took this magnitude to be the symmetric input, or the inhibition elicited by every stimulus regardless of direction^39^. Within this conceptual framework, null direction inhibition is the sum of directionally tuned and symmetric inputs. We thus took the directionally tuned component of inhibition to be the difference between null and preferred IPSC magnitudes (Fig. 2b,c insets). Symmetric inhibitory inputs onto ON DSGCs were strongly velocity tuned and increased rapidly for stimuli above 5 °/sec, while symmetric ON-OFF DSGC inhibition was only weakly tuned with respect to velocity (Fig. 2b,d, e). However, both cell types received directionally tuned inhibition that was largely untuned with respect to velocity (Fig. 2c,f,g). Thus ON DSGC velocity tuned inhibition appeared to be overlaid upon a substrate of velocity-invariant directional tuning also received by ON-OFF DSGCs. Note, directional spike tuning of ON DSGCs for high velocity RDKs seemed to be well explained by this mechanism, as IPSCs were directionally tuned for RDK motion (Extended Fig. 3).

Next, we measured excitatory postsynaptic currents (EPSCs) to assess contributions toward direction tuning across velocities. We analyzed the symmetric and directionally tuned components of excitation similarly to IPSCs, where symmetric excitation was taken to be the magnitude of null direction EPSCs and directionally tuned excitation was the difference in magnitude between preferred and null direction EPSCs. ON-OFF DSGC EPSCs were transient, and frequently directionally tuned (Fig. 3a). Interestingly, Trhr and Drd4 ON-OFF DSGC EPSCs, which prefer posterior motion, had a slight preference for higher velocities, whereas Hoxd10 ON-OFF DSGCs, which prefer anterior directed motion, had no velocity preference. These differences perhaps underlie a previously reported slight bias among Hoxd10 ON-OFF DSGCs for lower velocities relative to ON-OFF DSGCs^34^. In contrast, ON DSGC EPSCs had a large transient component, but also exhibited a sustained phase that lasted longer than ON-OFF DSGCs (Fig. 3a). Contrary to a previous report^27^, we saw minimal directional tuning in ON DSGC EPSCs (Fig. 3f). Similarly to IPSCs, we observed a minor EPSC OFF response in ON DSGCs. Because there was so little directionally tuned excitation for ON DSGCs, speed tuning indices were highly variable (Fig. 3d,g).

**Figure 3.**
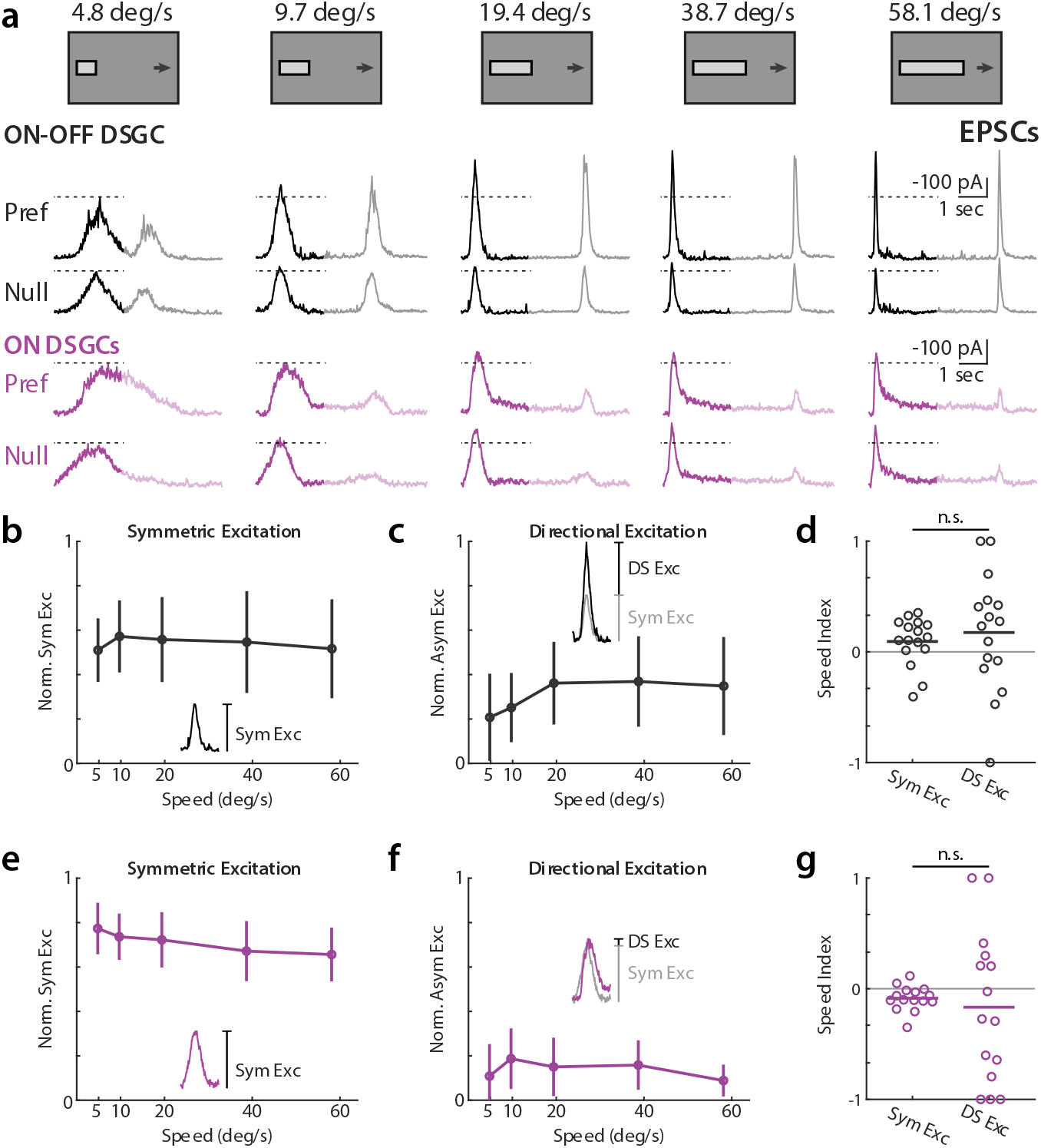
ON-OFF receive significant directionally tuned excitation while ON-DSGCs receive minimally directionally tuned excitation, both with weak velocity preference. (a) Example ON-OFF (*black*) and ON (*purple*) DSGC EPSC recordings for drifting bar stimuli, flipped vertically for display purposes. Opaque traces shows analysis window restricted to ON responses. Dashed lines indicate EPSC amplitude at the lowest tested velocity. (b) Population-averaged velocity tuning curves of symmetric excitation normalized to each cell’s maximal preferred direction EPSC. Error bars show standard deviation. Inset shows measurement of symmetric excitation as amplitude of null direction EPSC. (c) Population-averaged velocity tuning curves of directional excitation normalized to each cell’s maximal preferred direction EPSC. Error bars show standard deviation. Inset shows measurement of directional excitation as amplitude difference of preferred minus null direction EPSC. (d) Speed indices of symmetric and directional excitation. EPSC speed indices were compared via two-sided Wilcoxon signed-rank test; NS, *P* = 0.12, 16 cells in 10 mice. (e-g) Same as b-d, but for ON DSGCs. ON DSGC EPSC speed indices were compared via two-sided Wilcoxon signed-rank test; NS, *P* = 0.60, 15 cells in 10 mice.

We further investigated whether the time course of EPSCs and IPSCs contributed to DSGC tuning. We found that the peak amplitude and total charge transfer of EPSCs and IPSCs were well correlated, suggesting that the shape of postsynaptic currents did not provide significant additional explanatory power in describing DSGC spike tuning (Extended Fig. 4a,b). The relative DS indices of amplitude and total synaptic input can further illuminate the presynaptic origins of selectivity; direction selectivity that emerges from the coincident arrival of space-time wired inputs should be tuned in terms of event amplitude, but not charge transfer. We instead saw that the amplitude and charge transfer relation of synaptic events largely fell along unity, implying tuned synaptic release (Extended Fig. 4c,d). We also investigated the relative timing (measured as time of peak amplitude) between EPSCs and IPSCs as a potential source of DSGC tuning. On average, ON-OFF DSGC excitation led inhibition in the preferred direction, and lagged inhibition in the null direction, consistent with a ~50 μm preferred vs null spatial offset (Extended Fig. 5), similar to values reported elsewhere^42^. ON DSGCs excitation consistently lagged inhibition in both preferred and null directions, and thus seems unlikely to contribute to directional tuning (not shown). Thus, ON-OFF DSGCs appeared to use a combination of inhibitory, excitatory, and timing based mechanisms to support tuning, whereas ON DSGCs primarily relied on directional inhibition.

### Dynamic clamp experiments indicate synaptic inputs are sufficient to induce velocity tuning of DSGCs

Though voltage clamp experiments allow readout of synaptic conductances as measured at the soma, they do not allow for assessment of how different DSGC types integrate those inputs nor how that integration contributes to tuning for direction or velocity. Complex dendritic processing is known to occur in DSGCs^22,33,43^, but how integration properties might differentially influence temporal filtering properties of ON and ON-OFF DSGCs has not been studied. This question is further motivated by recent reports which suggest that parallel retinal ganglion cell circuits receiving similar synaptic inputs can craft distinct sensory representations due to differences in intrinsic biophysical properties^44,45^. To address this question, we used dynamic clamp to deliver synaptic inputs directly to the cell soma and test if distinct velocity tuning between DSGC types could be partially explained by differences in integrative properties. Using measured ON-OFF DSGC conductances, we delivered dynamic clamp inputs to ON and ON-OFF DSGCs and recorded spiking activity in the absence of visual input (Fig. 4a,b). We found that both ON and ON-OFF DSGC dynamic clamp driven activity resembled ON-OFF DSGC light driven spike firing. Velocity tuning for ON and ON-OFF DSGCs was similar when driven by ON-OFF DSGC dynamic clamp conductances (Fig. 4c,d). This suggests that our measured synaptic inputs are sufficient to explain the tuning of ON-OFF DSGCs, and that ON DSGC velocity tuning is not driven by differences in integrative properties.

**Figure 4.**
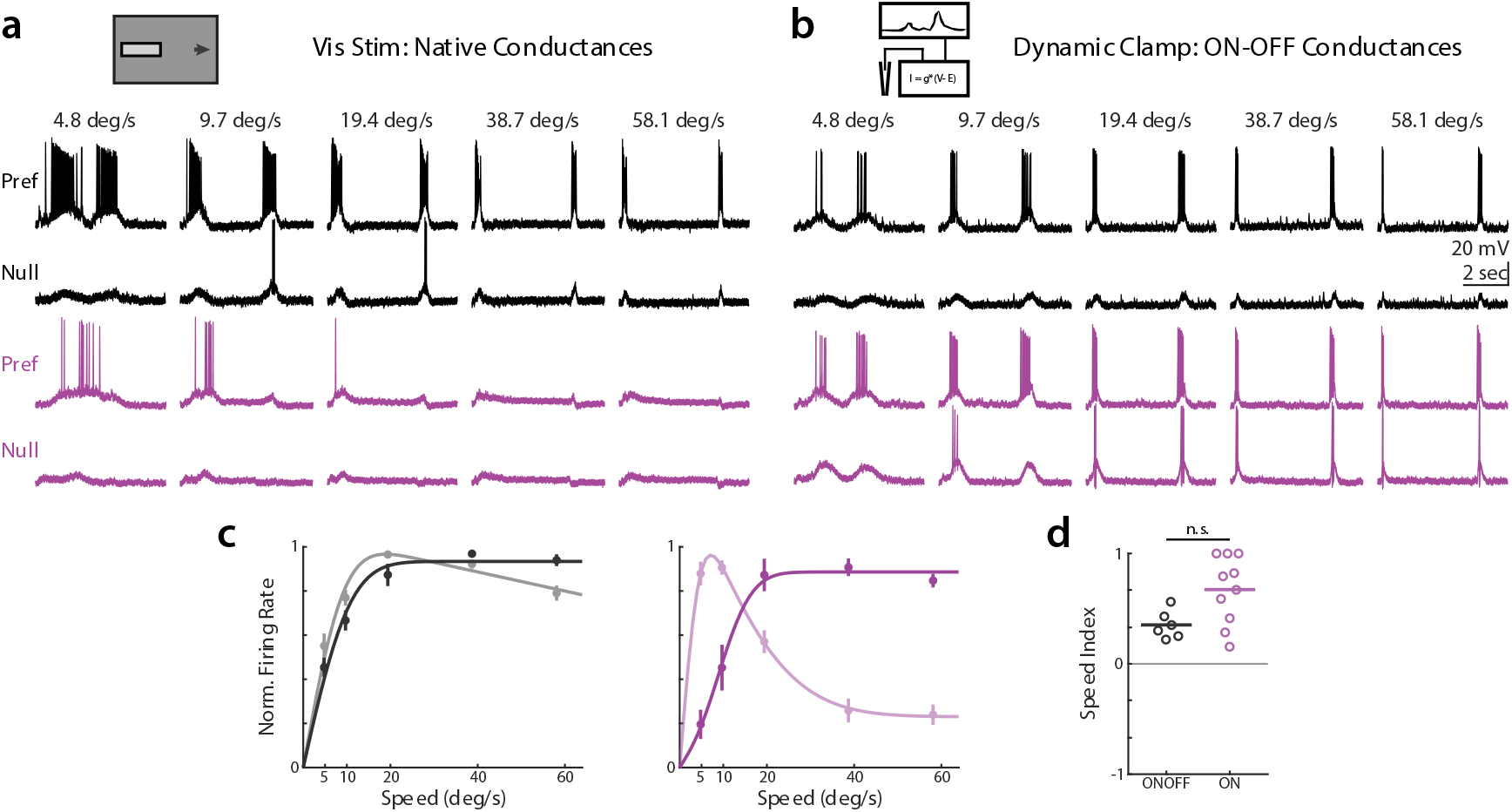
Dynamic clamp experiments indicate that integration and other biophysical properties of On-DSGCs do not dictate velocity tuning. (a) Example ON-OFF (*black*) and ON (*purple*) DSGC current clamp recordings for elongated bar visual stimuli where bar length scales with speed. (b) Same as a, but for stimulation with dynamic clamp inputs using ON-OFF DSGC conductances. (c) Population-averaged velocity tuning curves of dynamic clamp recordings normalized to peak firing rate. Error bars show standard error of the mean, solid lines show fits to an integral over difference of Gaussians. (d) Speed indices for ON-OFF and ON DSGCs stimulated via dynamic clamp. Comparison made via two-sided Wilcoxon rank-sum test; NS, *P* = 0.06, 6 ON-OFF DSGCs in 4 mice, 10 ON DSGCs in 6 mice.

### Conductance modeling indicates that asymmetric inhibition is the primary determinant of ON-OFF DSGC directional tuning

Given our measurements of excitatory, inhibitory, and timing computations all supporting ON-OFF DSGC directional selectivity, we wanted to assess the relative contribution of each of these mechanisms at each of our tested velocities. To assess the relative contributions of potential mechanisms for directional tuning, we used numerical modeling to simulate preferred and null direction depolarizations in a passive membrane model using the time-varying excitatory and inhibitory conductances we had recorded^46^ (Fig. 5a-c). Spiking is not included in this model, but is known to be nonlinear and enhance weak tuning that is already present^43,47^. By integrating our voltage clamp measured ON-OFF DSGC conductances in time, we were able to recapitulate the amplitude and directional selectivity of depolarizations measured in current clamp (Fig. 5d,e).

**Figure 5.**
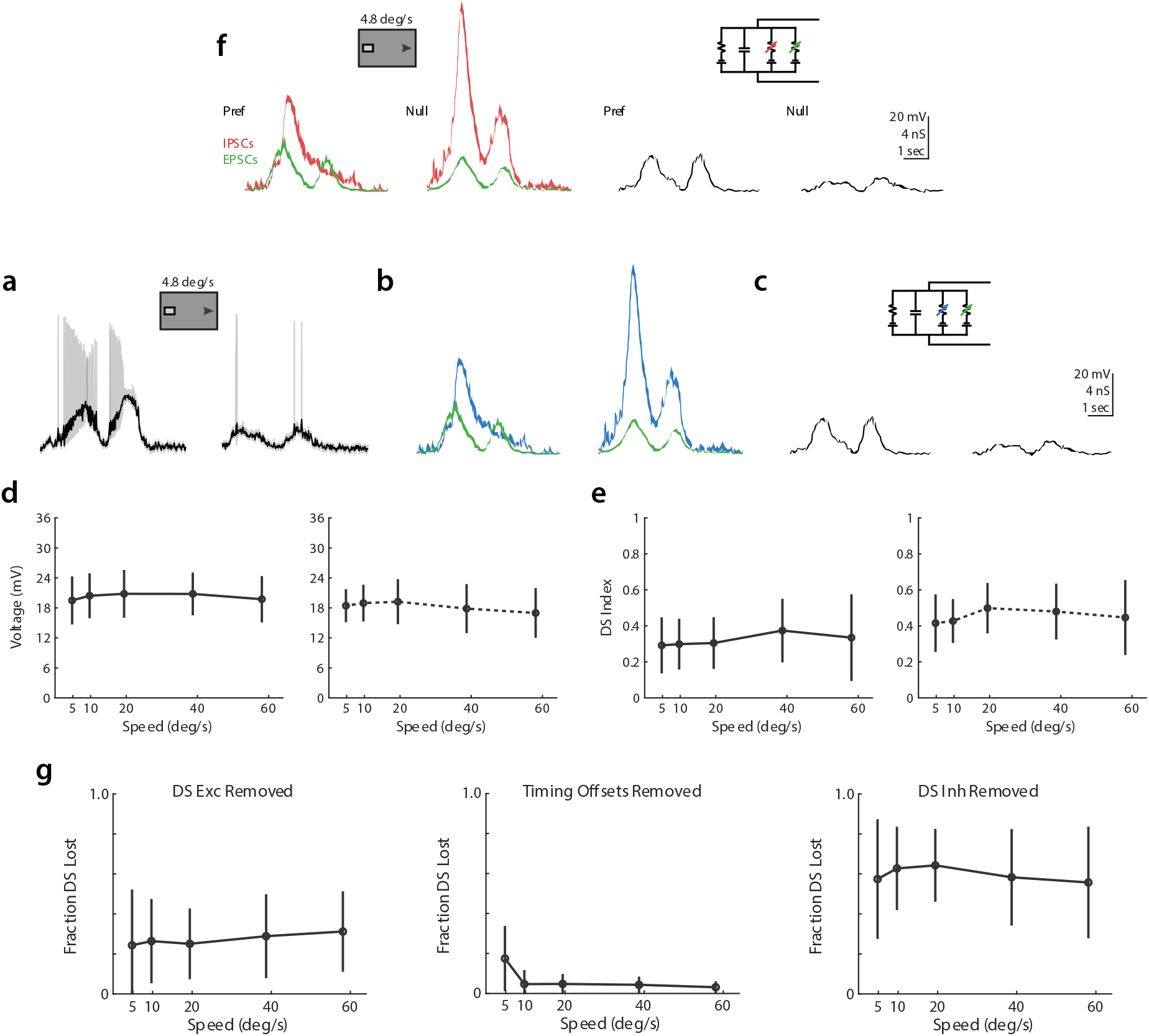
Conductance modeling indicates that asymmetric inhibition is the primary determinant of directional tuning. (a) Example preferred (*left*) and null (*right*) direction ON-OFF DSGC depolarizations, after removal of spikes via lowpass filtering. (b) Example EPSCs (*green, smaller*) and IPSCs (*blue, larger*) recorded from an ON-OFF DSGC in response to preferred (*left*) and null (*right*) bars as in a. (c) Example depolarizations from numerical integration of preferred (*left*) and null (*right*) conductances from b in a simple parallel conductance model, using forward Euler method. (d) Comparison of depolarizations measured in current clamp (*left*) and via conductance modeling (*right*). Markers show population averaged responses, and error bars show standard deviation. (e) Same as d, but for direction selectivity. (g) Velocity tuning of fractional direction selectivity loss for conductance model manipulations removing directional excitation (*left*), differential timing offsets (*middle*), and directional inhibition (*right*). Markers show population averaged responses, and error bars show standard deviation.

We then tested the contributions of excitatory, inhibitory, and timing based mechanisms by systematically removing the tuned components of each mechanism, integrating preferred and null direction depolarizations, and measuring the fractional loss of direction selectivity relative to our baseline models. Due to the shunting effects of inhibition, the sum total loss of direction selectivity from independently removing each mechanism is not necessarily expected to reach a 100% loss. Tuned excitation was removed by integrating a model where preferred direction excitatory conductances were substituted for those measured in the null direction, adding back in the appropriate time offset to recreate the relative timing of excitation and inhibition. With this manipulation, models experienced an average ~25% loss of direction selectivity, with minimal impact of velocity (Fig 5f). The impact of differential timing was assessed by integrating preferred and null conductances as normal, but shifting preferred direction inhibition forward in time to match the relative excitation and inhibition timing offset of the null direction. Models without a tuned timing mechanism experienced an average ~20% loss of direction selectivity at 5 °/sec, but a ~5% loss at higher velocities. Loss of differential timing had minimal impact on direction selectivity at higher velocities. Tuned inhibition was removed by substituting null direction inhibitory conductances for those measured in the preferred direction, again shifting in time to preserve the relative timing of excitation and inhibition. Loss of asymmetric inhibition resulted in an average ~60% reduction in direction selectivity, with minimal dependence on velocity. These models suggest that asymmetric inhibition is the primary driver of directionally tuned depolarizations, with asymmetric excitation playing a supplementary role and relative timing being a narrower contributor at low velocities.

### Direction and velocity tuned inhibition have different presynaptic origins

Finally, we sought to experimentally verify the independent origins of velocity and directionally tuned inhibition. To test whether reduction of directionally tuned inhibition was velocity dependent, we used *Vgat^flox/flox^/ Chat-IRES-Cre* mice to conditionally knock out vesicular GABA transporters in SACs and thereby prevent the release of GABA^31^. A previous study showed that this approach dramatically reduces directional tuning in posterior motion preferring ON-OFF DSGCs^31^, though the impact of this manipulation across velocities was not examined. We crossed *Vgat^flox/flox^/ Chat-IRES-Cre* with Hoxd10-GFP mice to assess the impact of SACs on the direction and velocity tuning of ON and anteriorly tuned ON-OFF DSGC inhibition. The initial transient component of ON-OFF DSGC IPSCs was remarkably diminished in these mice, as described previously^31^, leaving a sustained and directionally symmetric source of inhibition that persisted for the duration of the bar’s time within the receptive field (Fig. 6a). ON DSGC IPSCs retained a velocity tuned initial phasic response, though also lacked directional tuning (Fig. 6a). For both cell types, nearly all directionally tuned inhibition was abolished at every velocity (Fig. 6c,e). The velocity tuning of symmetric inhibition was similar between control and knockout animals, and we found no significant differences in speed tuning indices (Fig. 6b,d,f). This result is consistent with a previous study showing directionally tuned calcium transients in SAC varicosities at a range of velocities^26^.

**Figure 6.**
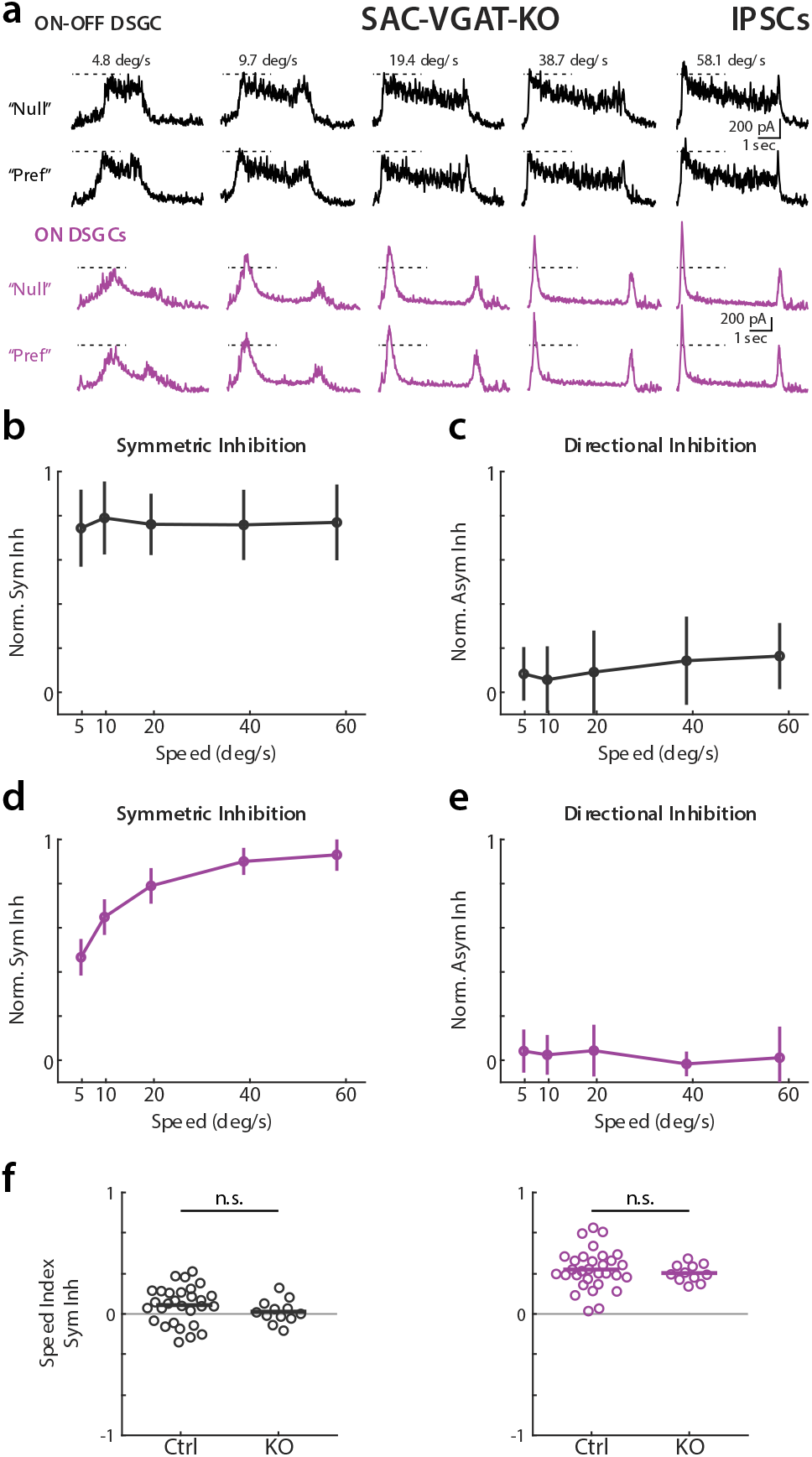
Elimination of GABA release from SACs reduces directional tuned inhibition across all velocities in On and ON-OFF DSGCs. a) Example ON-OFF (*black*) and ON (*purple*) DSGC IPSCs for elongated bar stimuli in *Hoxd10-GFP / Vgat^flox/flox^/ Chat-IRES-Cre* mice. Dashed lines indicate IPSC amplitude at the lowest tested velocity. (b) Population-averaged velocity tuning curves of symmetric inhibition in KO mice normalized to cell’s maximal IPSC. Error bars show standard deviation. (c) Population-averaged velocity tuning curves of directional inhibition normalized to cell’s maximal IPSC. (d-e) Same as b-c, but for ON DSGCs. (f) Symmetric inhibition speed indices in control and knockout animals. Comparison made via two-sided Wilcoxon rank-sum test; ON-OFF IPSCs, NS, *P* = 0.19, 29 cells in 19 control mice and 12 cells in 5 knockout animals. ON IPSCs, NS, *P* = 0.61, 31 cells in 21 control mice and 11 cells in 6 knockout animals

We next tested the impact of manipulating ON DSGC velocity preference on directional tuning. Previous work in the rabbit retina showed that ON DSGCs are inhibited by high temporal frequency flickering stimuli, and that glycine receptor antagonist strychnine blocks this frequency tuned inhibition^9,13^. We recorded ON DSGC IPSCs in response to elongated bar stimuli before and after bath application of strychnine. ON DSGC IPSCs remained directionally tuned after strychnine application, but showed a dramatic reduction in peak amplitude at higher velocities (Fig. 7a). Interestingly, OFF inhibition was often weakened but not completely abolished in ON DSGCs under strychnine application. Strychnine selectively abolished the velocity tuned component of ON DSGC symmetric inhibition, leaving IPSC magnitudes at the lowest velocities largely unchanged while substantially reducing inhibition for high velocity stimuli (Fig. 7b,d). There was no net effect of strychnine on the magnitude or velocity dependence of tuned inhibition, indicating selective disruption of velocity tuning pathways (Fig. 7c). ON DSGC current clamp recordings in the presence of strychnine were consistent with increased firing and stable direction selectivity for high velocity stimuli (Extended Fig. 6). We saw no effect of strychnine on ON-OFF DSGC IPSCs, though recent work suggests that glycinergic inhibition may play a role in SAC gain control and thus impact ON-OFF DSGC directional selectivity under certain stimulus conditions^48^. The data presented here suggest independent origins of symmetric and asymmetric inhibition, and that SACs are solely responsible for directional tuning independent of velocity in both ON and ON-OFF DSGCs.

**Figure 7.**
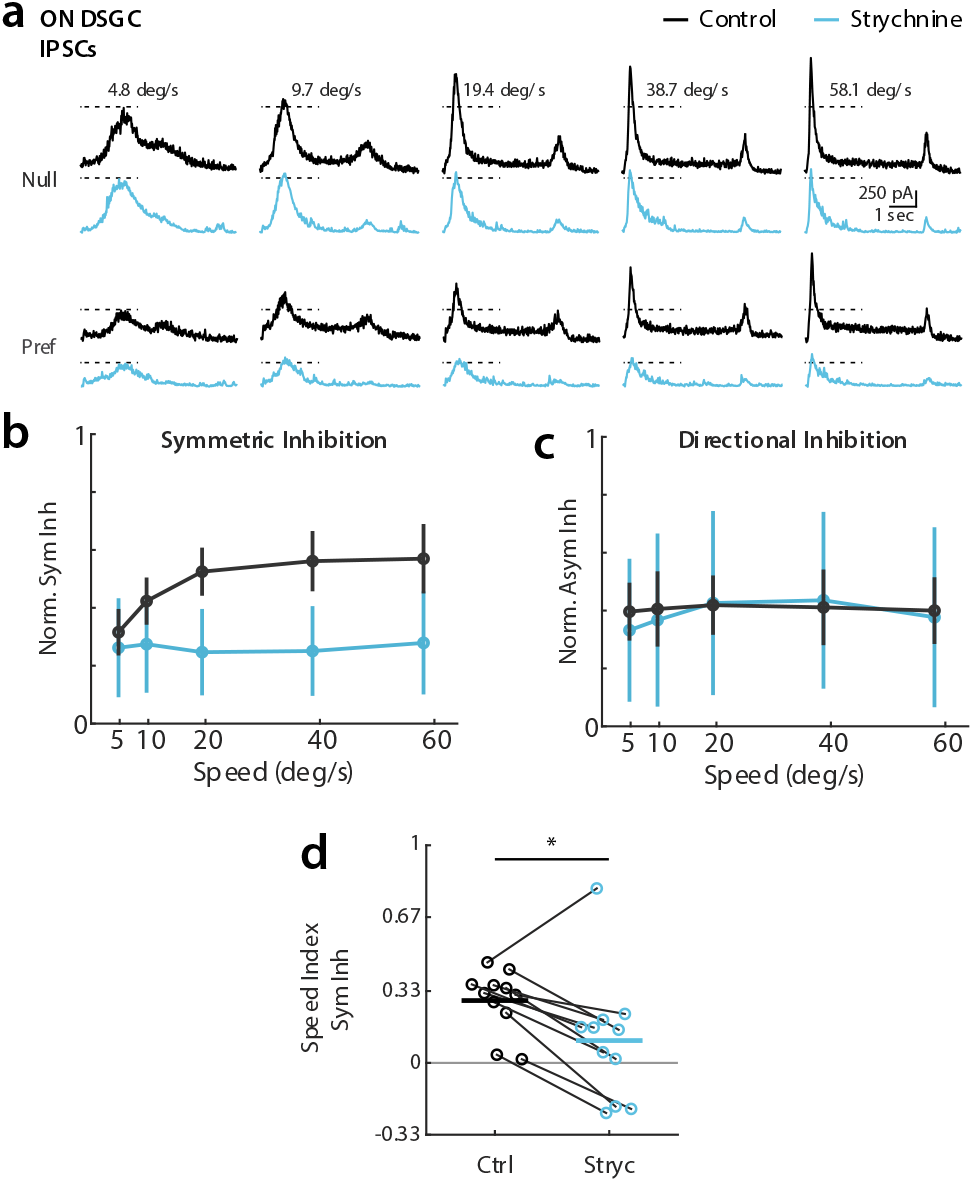
Blockade of glycine receptors eliminates the velocity tuned symmetric inhibition on ON-DSGCs. (a) Example ON DSGC IPSC recordings for elongated bar stimuli before (*black*) and after (*blue*) strychnine wash. Dashed lines indicate IPSC amplitude at the lowest tested velocity. (b) Population-averaged velocity tuning curves of symmetric inhibition for control and strychnine conditions normalized to cell’s maximal null direction IPSC before wash. Error bars show standard deviation. (c) Population-averaged velocity tuning curves of directional inhibition for control and strychnine conditions normalized to cell’s maximal null direction IPSC before wash. (d) Symmetric inhibition speed indices before and after strychnine wash. Comparison made via two-sided Wilcoxon signed-rank test; \**P* = 0.04, 11 cells in 8 mice.

## Discussion

The sensory periphery constructs distinct neural representations to subserve the goals of both reflexive and imaging forming vision^4^. Feature detectors in the periphery must efficiently compute representations of the external world, without the neuronal resources of central sensory brain regions. Here we demonstrate that distinct inhibitory pathways independently control tuning for the velocity and direction of motion on the retina. Namely, directionally tuned inhibition from starburst amacrine cells provides velocity invariant direction selective tuning to ON and ON-OFF DSGCs while symmetric inhibition from a glycinergic circuit provides directionally invariant velocity tuning to ON DSGCs. Dynamic clamp experiments show that synaptic inhibition is sufficient to explain differences in ON and ON-OFF DSGC velocity preference, and conductance modeling indicates that the impact of directional inhibition outweighs other tuning mechanisms in generating directionally tuned depolarizations. Thus, distinct inhibitory motifs emerge wherein the invariant motion feature tuning of amacrine cells form the computational building blocks of retinal outputs. Here we discuss the implications of these findings in the context of recent studies elucidating various circuit mechanisms that contribute to computations for direction and velocity.

### Synaptic origins of retinal direction selectivity

Asymmetric inhibition provided by SAC GABAergic synapses onto DSGCs is known to be a vital component of retinal direction selectivity^2,18,24^. SAC processes themselves are direction selective, releasing more GABA for outward motion from the soma. Thus, some elementary motion computations must occur within the SAC dendrite.

Our findings indicate that directionally tuned SAC inhibition to DSGCs is velocity invariant. First we show that the directionally tuned component of IPSCs remain constant in both ON and ON-OFF DSGCs across velocities (Fig. 2). Second, we show that elimination of GABA release from SACs abolishes directionally tuned inhibition at every tested velocity, and leaves behind non-directional, symmetric inhibition (Fig. 6). These findings are consistent with observations from 2-photon calcium imaging of SACs that outward motion preference is maintained in mice for velocities greater than ~3 °/sec^26^. However, the basis of this velocity invariance remains to be determined. In Ding et al, computational modeling was used to argue that reciprocal inhibition between SACs is a key driver of outward motion preference, and that the inter-soma distance between SACs determines the velocities at which direction selectivity is effectively computed. However, another study found that eliminating reciprocal inhibition via conditional knockout of SAC GABAA receptors had no effect upon ON pathway direction selectivity except during stimulation on top of inhomogeneous backgrounds^49^, and thus the true role of reciprocal SAC inhibition in velocity invariant computation perhaps remain unresolved. In contrast, other models of SAC tuning based on the spatial arrangement of different feedforward excitatory inputs with distinct kinetics imply velocity tuning in the SAC process^50^. Though the specific contributions of different mechanisms to SAC direction selectivity at different velocities is an area of ongoing research, our data shows explicitly for the first time that SAC based inhibition to both ON and ON-OFF DSGCs is directionally tuned independent of velocity.

Other mechanisms in addition to asymmetric inhibition have been implicated in contributing to DSGC directional tuning. Cholinergic excitation from SACs is known to contribute to ON-OFF DSGC depolarizations, and may be responsible for establishing timing offsets between excitation and inhibition^31,42,51^. Growing evidence from several recent studies suggests that motion tuning is also present within the synaptic boutons of many bipolar cells^52–54^. Interestingly, directional tuning in bipolar cells presynaptic to ON-OFF DSGCs appears to be inherited from SACs, perhaps explaining the weak velocity dependence of directional excitation observed in our study^52,53^. This link highlights the value of conductance modeling in allowing for independent interrogation of the impact of excitation and inhibition on directional tuning (Fig. 5), whereas experimental manipulations may struggle to disentangle these mechanisms. Another study described directional excitation as an important source of ON DSGC tuning^27^. We observed directional excitation in ON-OFF DGCs, but saw minimal tuned excitation in ON DSGCs; whether this difference was due to differences in ON DSGC subtypes, stimulus parameters, or other recording conditions may require additional investigation.

What is the utility in having multiple mechanisms contributing to DSGC directional? Our data points to directional inhibition as the dominant contributor to directional tuning across physiological velocities, and argues against a combination of mechanisms upholding computation at distinct speeds. One possible role for supplementary mechanisms is to support computation across spatial scales. Integration of various mechanisms at the local dendritic level may enable reliable directional discrimination for small stimuli within subsets of the ON-OFF DSGC receptive field^33,55^. This hypothesis is supported by our observations that ON DSGCs, which are thought to be global motion detectors, generally lacked additional tuning mechanisms beyond directional inhibition. Additional mechanisms may also play important roles under more naturalistic, complex stimulus conditions, or may contribute to ON-OFF DSGC tuning for pattern versus component motion^11,49,56^.

### Symmetric inhibition shapes spatial and temporal tuning properties of DSGCs

We provide several lines of evidence that ON DSGC preference for low velocity motion is mediated by symmetric, non-directional inhibition. Firstly, we show that symmetric inhibition is tuned for high velocities (Fig. 2). Secondly, dynamic clamp conductances are sufficient to recapitulate ON-OFF DSGC velocity tuning, arguing against cell intrinsic differences shaping velocity preference (Fig. 4). Finally, strychnine reduces ON DSGC velocity tuning, leaving directionally tuned but velocity invariant IPSCs (Fig. 7). Contrary to classical models of elementary direction computation, these data support a model wherein velocity and direction tuning are each independently conferred by distinct sources.

Glycinergic amacrine cells form synaptic connections throughout the inner retina, with targets including bipolar cell terminals, retinal ganglion cells, and other amacrine cells^57^. A recent study has also implicated glycinergic circuitry in SAC gain control^48^. Our data is consistent with a feedforward glycinergic circuit providing symmetric, velocity tuned inhibition to ON but not ON-OFF DSGCs, similar to a circuit previously described in the rabbit retina that suppresses ON DSGCs spiking to rapid saccade-like stimuli^9^. A recent study implicates VGlut3 amacrine cells, which were previously thought to provide solely excitatory input to ON DSGCs^27,58,59^, as being a possible source of this velocity tuned glycinergic inhibition^60^.

This finding of a non-directional inhibitory source of ON DSGC velocity tuning is in contrast to excitation based correlation type models that jointly encode direction and velocity^27,61^. These space-time wiring models rely on the alignment of excitatory inputs with distinct kinetics such that inputs are differentially summed in preferred but not null directions, even though the total synaptic input is not itself tuned. We saw minimal evidence of tuned excitation in ON DSGCs, and the directional tuning of the amplitude and charge transfer of ON-OFF DSGC excitation was well correlated (Extended Fig. 4), contrary to the predictions of these models. Further, successful directional computation within these space-time wiring models should be highly dependent upon the spatial and temporal frequencies of the stimulus. We found that our RDK stimuli, which contain a broad spectrum of spatiotemporal frequencies due to the uneven spacing of individual dots, revealed velocity invariant directional tuning in ON DSGCs (Fig. 1). We postulate that this is due to weak activation of the glycinergic input that mediates inhibition for full field motion^60^.

We also found ON-OFF DSGCs possess a symmetric source of inhibition that is GABAergic and persists in the absence of GABA release from SACs (Fig. 6). This symmetric inhibition has also been reported in posteriorly tuned ON-OFF DSGCs^31,39^. One potential source of this inhibition is the VIP+ amacrine cell, which is known to synapse onto ON-OFF DSGCs, but whose role in shaping DSGC activity remains mysterious^40^. Interestingly, inhibition from spiking wide field amacrine cells has been shown to mediate size tuning in DSGCs via presynaptic inhibition^15^. Though not directly targeting retinal ganglion cells, this circuit provides a clear parallel to our findings of distinct inhibitory motifs for direction and velocity tuning. These results suggest an elegant degree of parallel computation within the retina, where invariant representations of visual features are constructed by inhibitory interneurons to subserve the feature tuning of diverse retinal ganglion cells.

## Methods

### Animals

All animal procedures were approved by the UC Berkeley Institutional Animal Care and Use Committee and conformed to the NIH Guide for the Care and Use of Laboratory Animals, the Public Health Service Policy, and the SFN Policy on the Use of Animals in Neuroscience Research. Retinas from adult mice (P28-P80) of either sex were prepared as previously described^62^, but in brief were dissected under infrared illumination, mounted over a 1-2 mm^2^ hole in filter paper, and stored in oxygenated Ames’ media in the dark at room temperature. ON DSGCs were targeted for current clamp, voltage clamp, and dynamic clamp experiments in Hoxd10-GFP (Tg(Hoxd10-EGFP)LT174Gsat/Mmucd) animals^34^, which were sometimes crossed with *Vgat*^flox/flox^ (*Slc32a1<tm1Lowl>/J*) or *ChAT-IRES-Cre* (129S6-*Chat*^tm2(cre)Lowl^/J) mice. ON-OFF DSGCs were targeted in these mice, and additionally in Trhr-GFP (B6;FVB-Tg(Trhr-EGFP)HU193Gsat/Mmucd) and Drd4-GFP (Tg(Drd4-EGFP)W18Gsat/Mmnc) mice^12,35^. Similar responses were observed for cells recorded from each of these different transgenic lines. Knockout experiments were performed on Hoxd10 / *Vgat*^flox/flox^ / *ChAT-IRES-Cre* mice, which were generated by crossing each line. ON and ON-OFF DSGCs were distinguished in Hoxd10 lines on the basis of response polarity to a brief light step and morphological stratification.

### Two-photon targeted whole-cell recordings

Retinas were placed under the microscope in oxygenated Ames' medium at 32– 34°C. GFP+ cells were identified using a two-photon microscope tuned to 920 nm to minimize bleaching of photoreceptors. The inner limiting membrane above the targeted cell was dissected using a glass electrode. Current clamp and dynamic clamp recordings were conducted with internal solution composed, in mM, of: 115 K+ gluconate, 9.7 KCl, 1 MgCl2, 0.5 CaCl2, 1.5 EGTA, 10 HEPES, 4 ATP-Mg2, 0.5 GTP-Na3, 0.025 TexasRed (pH = 7.2 with KOH, osmolarity = 290). Voltage clamp recordings were performed with internal solution containing the following, in mM: 110 CsMeSO4, 2.8 NaCl, 20 HEPES, 4 EGTA, 5 TEA-Cl, 4 Mg-ATP, 0.3 Na3GTP, 10 Na2Phosphocreatine, 5 QX-Br, 0.025 Texas Red (pH = 7.2 with CsOH, osmolarity = 290). Holding voltages for measuring excitation and inhibition after correction for the liquid junction potential (−10 mV) were −60 mV and 0 mV, respectively. Signals were acquired using pCLAMP 9 recording software and a Multiclamp 700A amplifier (Molecular Devices), sampled at 10 kHz, and low-pass filtered at 6 kHz. Strychnine experiments were performed in Ames’ media with 2 μM strychnine hydrochloride (Sigma-Aldrich).

### Visual stimuli

Visual stimuli were generated via custom MATLAB functions written with Psychophysics Toolbox on a computer running a 60 Hz DMD projector (EKB Technologies) with a 485 nm LED light source. The DMD image was projected through a condenser lens, and aligned on each experimental day to the photoreceptor layer of the sample. All stimulus protocols were centered on the soma of the recorded cell, and were presented after at least 10 seconds of adaptation on a dark background (9.4 × 10^3^ R*/rod/s). Bars, gratings, and dots were all of positive contrast, and equal intensity (2.6 × 10^5^ R*/rod/s).

Baseline direction selectivity was first assessed with 100 μm (3.2 °) wide by 650 μm (21 °) long bars moving at either 200 μm/s or 500 μm/s (6.5 °/s and 16.1 °/s respectively). Responses were recorded for at least 3 repetitions of bars moving in 8 block shuffled directions, each separated by 45 degrees. Online analysis was then used to determine a DSGC’s preferred direction for velocity stimuli: for current clamp, this was the vector sum angle of spike counts, for voltage clamp, this was 180 degrees offset from the vector sum angle of IPSC magnitudes. For voltage clamp experiments on knockout mice lacking directional inhibition, two orthogonal “preferred” and “null” axes were used for velocity experiments to ensure at least four total directions were probed for residual tuning.

Elongated moving bars and drifting gratings were presented for at least 3 repetitions in preferred and null directions at block shuffled velocities. Moving bars were 100 μm (3.2°) wide and ranged from 150 – 1800 μm/s (4.8 – 58.1 °/s). Drifting gratings were presented for 6 seconds each within a 300 μm radius mask, and had a 250 μm spatial period. Temporal frequencies were varied from 0.2 to 7.2 cycles/sec (1.6 – 58.1 °/s).

Random dot kinetogram (RDK) stimuli matched previously described parameters^38^. Direction selectivity in response to RDK motion was assessed similarly to baseline measurements, with at least 3 repetitions of RDKs moving in 8 block shuffled directions. RDK stimuli were presented for 5 seconds each at 20% density and were 100% coherent. Individual dots were 62 μm (2 °) in diameter and moved at 775 μm/s (25 °/s).

### Dynamic clamp

We constructed a microcontroller based dynamic clamp device following published specifications^63^, also found on dynamicclamp.com. The current (*I*) delivered to a cell was calculated as:

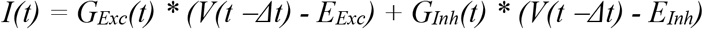

Where *G_Exc_* and *G_Inh_* are the respective time varying conductance traces for excitation and inhibition recorded from elongated bar visual stimuli, *V* is the cell membrane potential, and *E_Exc_* and *E_Inh_* are 0 mV and −60 mV reversal potentials respectively. Conductances used as dynamic clamp inputs were taken from individual cells and were averaged over 3 trials for each velocity of visual stimulus. At least 3 repetitions of preferred and null direction conductances were presented for dynamic clamp experiments, at block shuffled velocities.

### Conductance modeling

The contributions of synaptic conductances to tuned depolarizations were simulated via a parallel conductance model implemented in MATLAB^46^. Conductances *G_Exc_* and *G_Inh_* used as model inputs were taken from individual cell voltage clamp recordings in response to elongated drifting bars, and were rectified and trial averaged for each velocity of visual stimulus. For each velocity, a simulated cell’s voltage time series trace was numerically integrated via the forward Euler method:

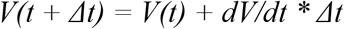

Where *dV / dt* was derived from the current flow across an RC circuit with empirically determined values for capacitance *C_m_* (80 pF), resting conductance *G_Leak_* (4.2 nS) and resting membrane potential *E_Leak_* (−55 mV):

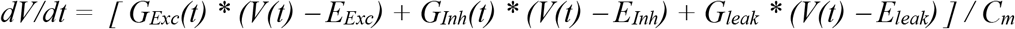

The amplitude of simulated depolarizations was compared between preferred and null directions, and direction selectivity indices were calculated at each speed.

Manipulations of specific tuning mechanisms were made by swapping or shifting in time the conductances used to integrate voltage. In each case, fractional loss of directional selectivity was assessed for a simulated cell at a given velocity via:

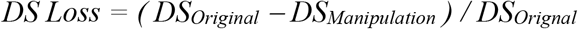

Where *DS_Original_* and *DS_Manipulation_* are the direction selectivity indices (see below) of a simulated cell at a given velocity.

The impact of three model manipulations was assessed. (1) Removal of asymmetric excitation was simulated by integrating null direction depolarizations as normal, but substituting in null for preferred direction excitation when integrating preferred direction depolarizations. This null swapped excitation was appropriately shifted in time so as to preserve the same preferred direction timing offset between peak excitation and inhibition. (2) Removal of differential timing offsets was simulated by integrating null direction depolarizations as normal, but shifting preferred direction inhibition (almost exclusively forward) in time to match the excitation and inhibition timing offsets measured in the null direction. (3) Removal of asymmetric inhibition was simulated by integrating preferred direction depolarizations as normal, but substituting in preferred for null direction inhibition when integrating null direction depolarizations. This preferred swapped inhibition was appropriately shifted in time so as to preserve the same null direction timing offset between peak excitation and inhibition.

### Analysis

Analysis was performed using custom MATLAB scripts. All analyses of drifting bar stimuli were restricted to the ON window immediately subsequent to a bar entering the DSGC’s receptive field. Grating and RDK analyses utilized the full period for which a stimulus was present.

Instantaneous firing rates were determined from current clamp and dynamic clamp data via kernel density estimation with a 200 ms Gaussian kernel. Peak firing rate was then taken to be the maximal instantaneous firing rate achieved within the analysis window. Voltage clamp data was baseline subtracted and lowpass filtered at 30 Hz. Peak current amplitudes and total charge transfer were calculated within the aforementioned analysis window. Depolarizations were measured from current clamp data by removing spiking activity via lowpass filtering at 20 Hz, and then measuring amplitude by baseline subtraction.

Directional selectivity indices were calculated from responses to preferred (*R_Pref_*) and null (*R_Null_*) direction motion as:

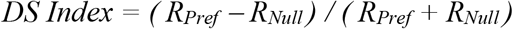

For current clamp and dynamic clamp recordings, responses were measured from peak firing rate. Peak EPSC and IPSC magnitudes were used for voltage clamp recordings. The signs of DSIs calculated from IPSCs were flipped to better reflect their contributions toward tuning. For conductance model simulated and lowpass filtered current clamp depolarizations, responses were measured as the depolarization amplitude from baseline. Negative values (rare cases where *R_Null_* > *R_Pref_*) where rectified to zero.

Speed indices were calculated from preferred direction responses to high (*R_High_*) and low (*R_Low_*) velocity motion as:

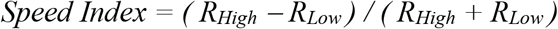

Values thus tended toward −1 for responses tuned to low velocities and toward +1 for responses tuned to high velocities, while zero indicated equal responses to high and low speeds. Due to the lengthy recordings required to isolate ON responses for low velocity object motion, *R_Low_* was determined from responses to 4.8 °/s moving bars, while 1.6 °/s was used for analysis of drifting grating responses. Responses to 58.1 °/s motion were used for *R_High_* in both gratings and bars.

### Statistics

Details of statistical tests, number of replicates, and p values are indicated in the figures and figure captions. Statistical methods were not used to predetermine sample size.

## Code and data availability

Modeling data and code can be found at https://github.com/FellerLabCodeShare/DSGC-Velocity-Project. Additional datasets and analysis code are available upon request.

## Acknowledgments

M.T.S. was supported by the National Science Foundation Graduate Research Fellowship (DGE 1752814). M.T.S. and M.B.F. were supported by NIH grants R01EY019498, R01EY013528, and P30EY003176.

## Extended Figures

**Extended Data Table 1.** Speed tuning indices. Speed tuning indices of DSGC inputs and outputs. Index values range from −1, which indicates preference for low velocities, to +1, which indicates preference for high velocities. Values tending toward zero indicate equal responses to high and low velocities. All P values are comparisons made to a zero median distribution via Wilcoxon signed-rank test.

| Cell Type | Measure | Condition | Mean | Std Dev | P Value | Cells | Mice |
| --- | --- | --- | --- | --- | --- | --- | --- |
| ON-OFF | Spikes | Bars | 0.21 | 0.23 | $3.8 \times 10^{-3}$ | 16 | 11 |
| ON-OFF | Spikes | Gratings | 0.54 | 0.32 | 0.01 | 8 | 5 |
| ON | Spikes | Bars | -0.63 | 0.28 | $8.4 \times 10^{-5}$ | 21 | 17 |
| ON | Spikes | Gratings | -0.91 | 0.25 | 0.02 | 7 | 6 |
| ON-OFF | Sym. Inh. | Bars | 0.07 | 0.15 | 0.04 | 29 | 19 |
| ON-OFF | DS Inh. | Bars | 0.06 | 0.21 | 0.05 | 29 | 19 |
| ON-OFF | Sym. Exc. | Bars | 0.09 | 0.22 | 0.09 | 16 | 10 |
| ON-OFF | DS Exc. | Bars | 0.18 | 0.53 | 0.18 | 16 | 10 |
| ON | Sym. Inh. | Bars | 0.36 | 0.16 | $1.2 \times 10^{-6}$ | 31 | 22 |
| ON | DS Inh. | Bars | -0.13 | 0.21 | $3.1 \times 10^{-3}$ | 31 | 22 |
| ON | Sym. Exc. | Bars | -0.09 | 0.11 | 0.01 | 15 | 10 |
| ON | DS Exc. | Bars | -0.17 | 0.68 | 0.39 | 15 | 10 |
| ON | Sym. Inh. | Bars, Strychnine | 0.10 | 0.29 | 0.52 | 11 | 8 |
| ON-OFF | Sym. Inh. | Bars, VGAT KO | 0.02 | 0.10 | 0.62 | 12 | 5 |
| ON | Sym. Inh. | Bars, VGAT KO | 0.34 | 0.07 | $9.7 \times 10^{-4}$ | 11 | 6 |
| ON-OFF | Spikes | Dynamic Clamp | 0.35 | 0.13 | 0.03 | 6 | 4 |
| ON | Spikes | Dynamic Clamp | 0.67 | 0.31 | $2.0 \times 10^{-3}$ | 10 | 6 |

**Extended Figure 1.**
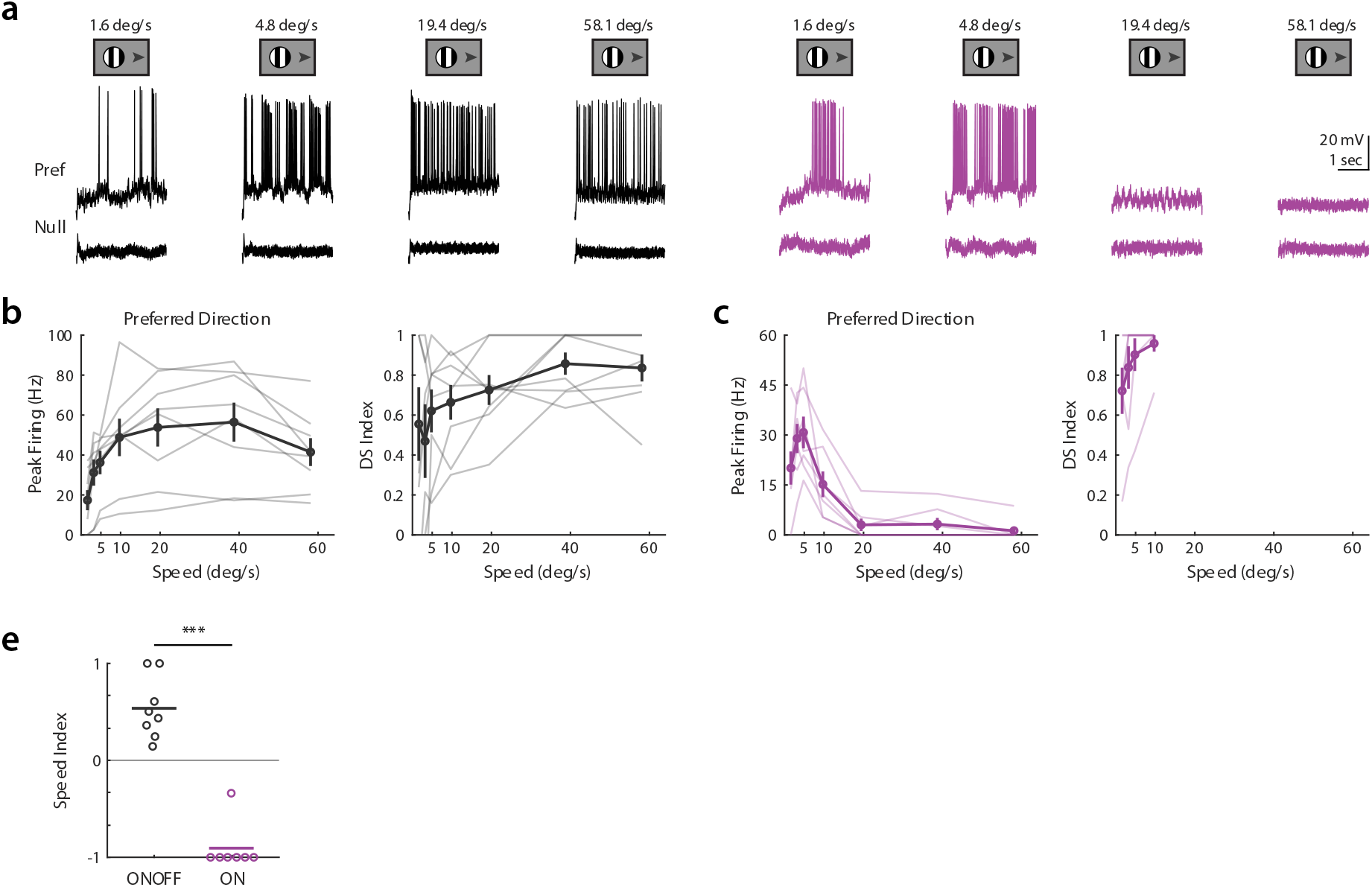
Velocity tuning of DSGCs not altered by using drifting gratings. (a) Example current clamp recordings *(ON-OFF black, ON purple*) in response to gratings drifting at several temporal frequencies in the cell’s preferred or null direction. (b) ON-OFF DSGC dependence of preferred direction spiking (*left*) and directional selectivity (*right*) on grating velocity. (c) Same as in b, but for ON DSGCs. Directional selectivity is difficult to assess at high velocities due to low overall spiking. (e) Speed tuning indices of preferred direction spiking. Comparison made via two-sided Wilcoxon rank-sum test; \*\*\**P* = 3.1 × 10^−4^, 8 ON-OFF DSGCs in 5 mice, 7 ON DSGCs in 6 mice.

**Extended Figure 2.**
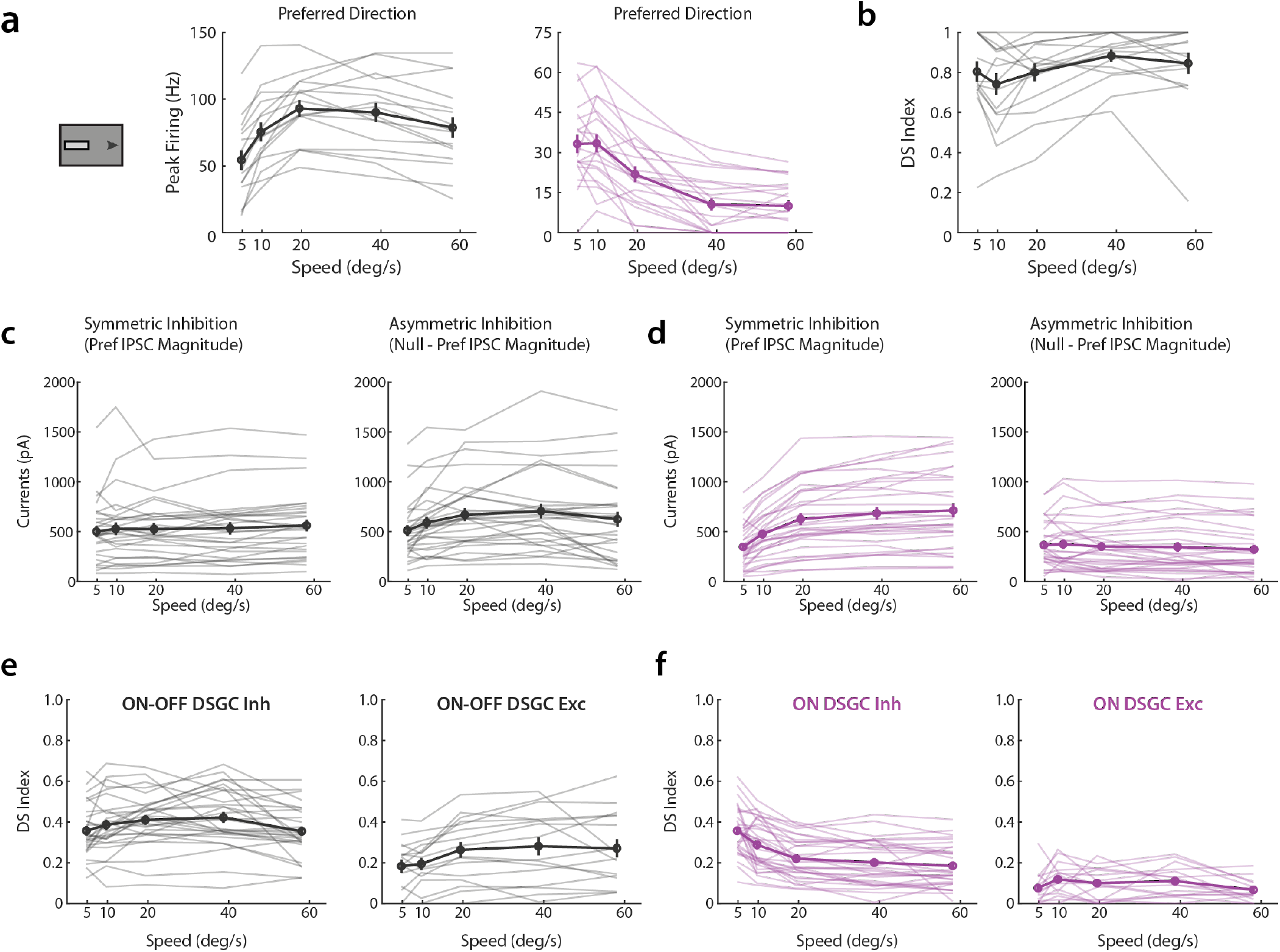
Data for Figures 1–3 presented as unnormalized values. (a) Summary plot of unnormalized velocity tuning curves based on current clamp recordings of ON-OFF (*black*) and ON DSGCs (*purple*) in response to drifting bar stimuli. In this and all other panels, transparent lines show individual cells while bold lines are the population average. Error bars show standard error of the mean. (b) ON-OFF DSGC direction selectivity indices versus velocity. ON DSGC directional selectivity is difficult to assess at high velocities due to low overall spiking and so is not included. (c) Summary plot of unnormalized velocity tuning curves of ON-OFF DSGC symmetric (*left*) and asymmetric (*right*) inhibition based on voltage clamp recordings. Symmetric inhibition was defined as the amplitude of preferred direction IPSCs, while asymmetric inhibition was measured as the amplitude of null direction IPSCs minus the magnitude of symmetric inhibition. (d) Same as in c, but for ON DSGCs. (e) ON-OFF DSGC direction selectivity indices of synaptic inputs versus velocity. Direction selectivity indices calculated directly from preferred versus null IPSC (*left*) and EPSC (*right*) amplitudes. (f) Same as e, but for ON DSGCs.

**Extended Figure 3.**
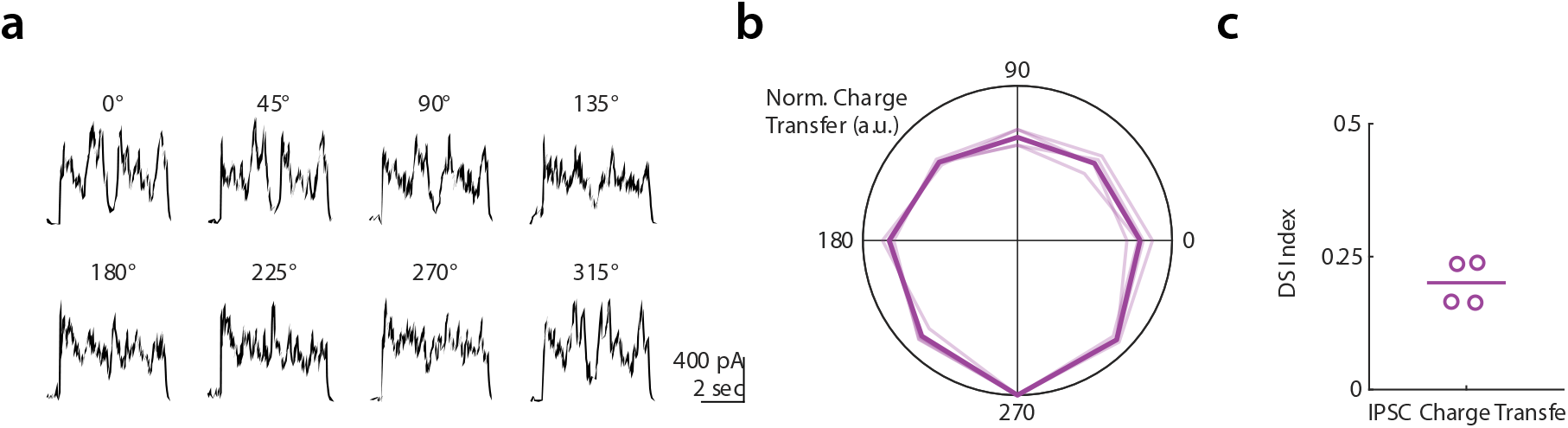
Voltage clamp recordings from ON DSGCs in response to random dot kinetogram show directional tuning. (a) Example ON DSGC inhibitory postsynaptic currents (IPSCs) for random dot kinteogram (RDK) stimuli moving coherently in one of eight directions. (b) Polar plot directional tuning curves of ON DSGCs normalized inhibitory charge transfer for RDK stimuli. Directions of maximal inhibition are aligned to 270 degrees. Transparent lines are tuning of individual cells, bold line is population average. (c) Direction selective indices computed from inhibitory charge transfer.

**Extended Figure 4.**
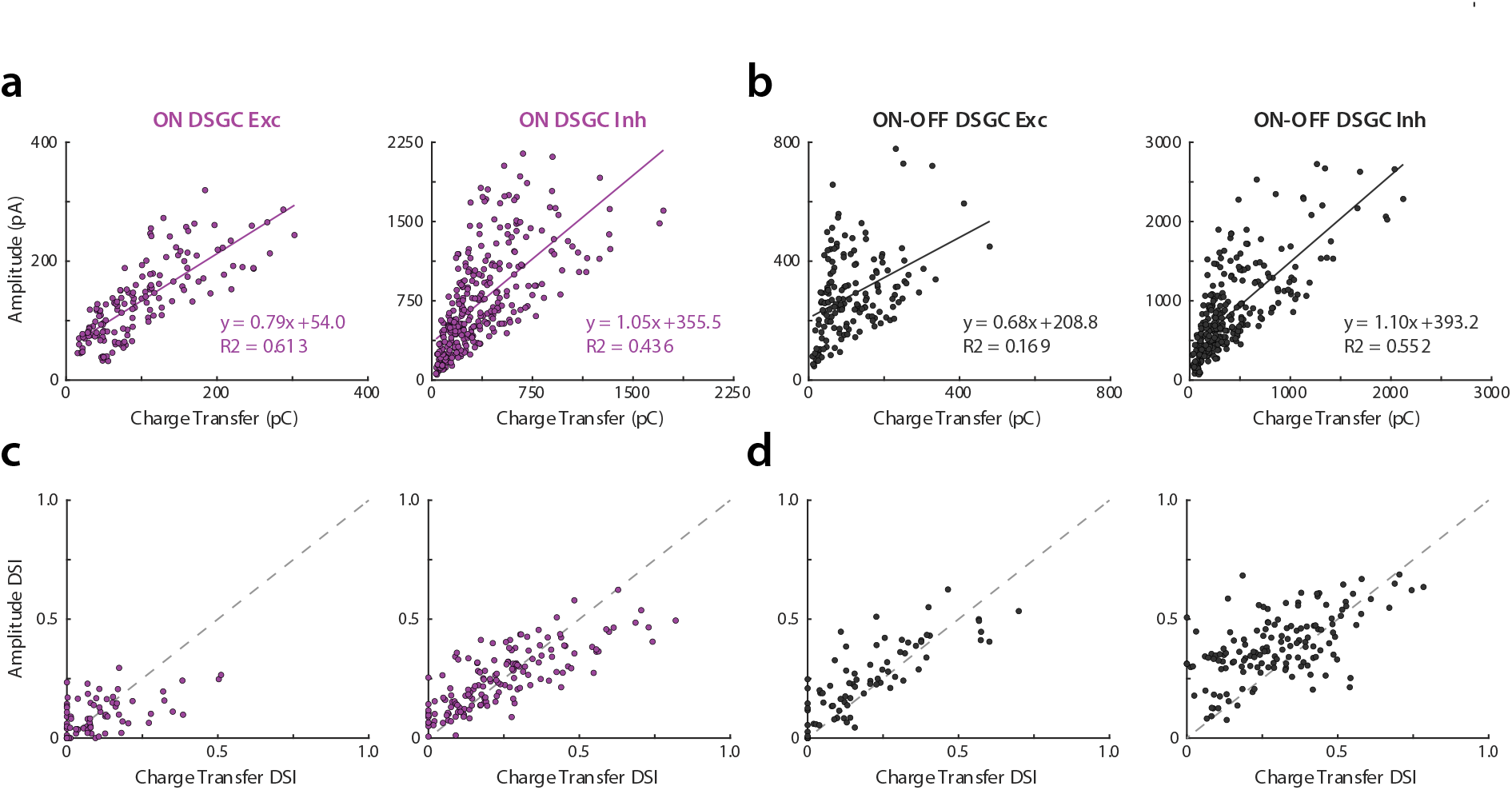
Total synaptic input is well approximated by EPSC and IPSC amplitudes. (a) Relationship between ON DSGC EPSC (*left*) / IPSC (*right*) amplitude and charge transfer. Individual points are the mean values for a given cell at a set velocity. Solid lines are linear fits, with slope, intercept, and coefficient of determination inset. (b) Same as a, but for ON-OFF DSGCs. (c) Relationship between ON DSGC direction selectivity of EPSC (*left*) / IPSC (*right*) amplitude and charge transfer. Individual points are the value for a given cell at a set velocity. Dashed gray line is unity. (d) Same as c, but for ON-OFF DSGCs.

**Extended Figure 5.**
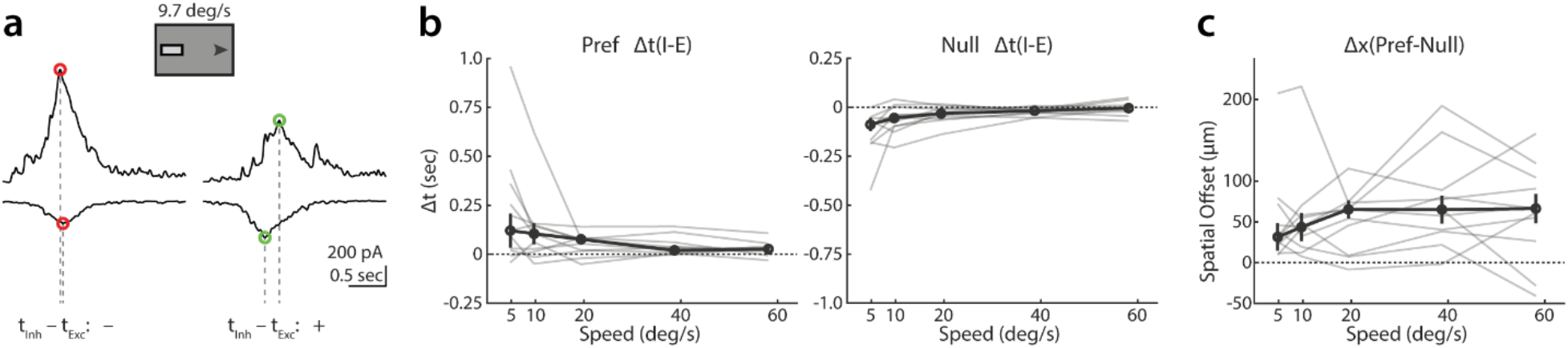
Method for determining the relative timing differences between excitation and inhibition. (a) Example ON-OFF DSGC IPSCs (*top*) and EPSCs (*bottom*) illustrating relative timing differences for preferred (*right*) and null (*left*) directed stimuli. IPSC peaks preceding EPSC peaks are treated as negative timing differences, while EPSCs preceding IPSCs are treated as positive timing differences. (b) Dependence of ON-OFF DSGC I-E timing differences on velocity for preferred (*left*) and null (*right*) directed stimuli. Transparent lines show individual cells, bold line is population average. Error bars show standard error of the mean. (c) Directionally tuned component of timing differences represented as a spatial offset. Transparent lines show individual cells, bold line is population average. Error bars show standard error of the mean.

**Extended Figure 6.**
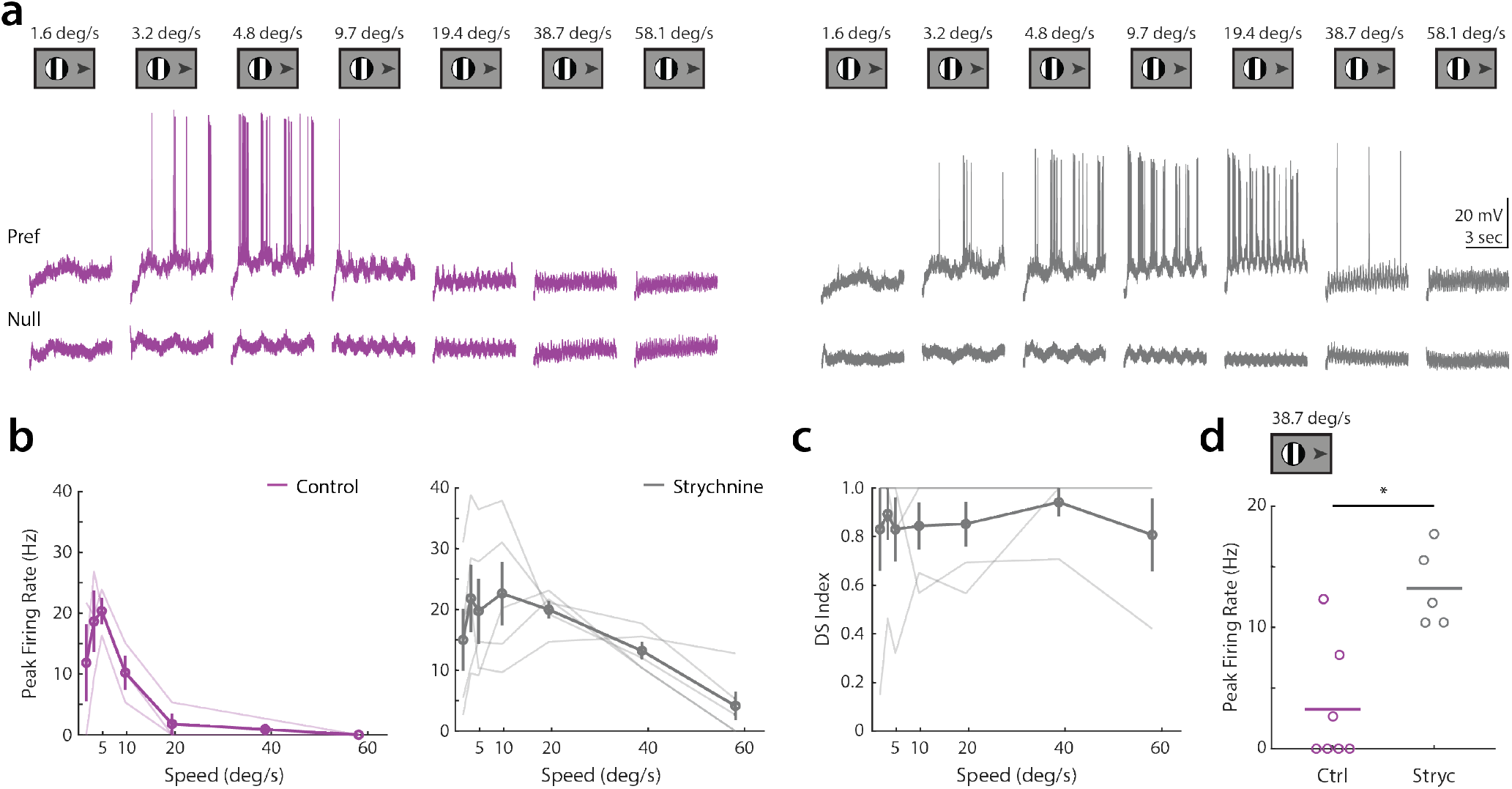
Glycine receptor antagonist strychnine increases ON DSGC spiking at high velocities. (a) Example ON DSGC current clamp recordings for gratings drifting at several temporal frequencies in the cell’s preferred or null direction, both before (*purple, left*) and after (*gray, right*) strychnine wash. (b) Dependence of preferred direction peak firing rate on drifting grating velocity, before (*left*) and after (*right*) strychnine wash. Transparent lines show individual cells, bold line is population average. Error bars show standard error of the mean. (c) Dependence of direction selectivity index on drifting grating velocity after strychnine wash. Transparent lines show individual cells, bold line is population average. Error bars show standard error of the mean. (d) Comparison of peak preferred direction firing rate at the second highest tested velocity (38.7 deg/sec) in control and strychnine conditions. Lines indicate cells for which recordings both before and after wash were collected. Comparison made via two-sided Wilcoxon rank-sum test; \**P* = 0.02, control 7 cells in 6 mice, strychnine 5 cells in 3 mice.

## References

1. Baden, T., Berens, P., Franke, K., Román Rosón, M., Bethge, M., and Euler, T. (2016). The functional diversity of retinal ganglion cells in the mouse. Nature 529, 345–350.

2. Summers, M.T., El Quessny, M., and Feller, M.B. (2021). Retinal Mechanisms for Motion Detection. Oxford Res. Encycl. Neurosci., 1–24.

3. Schwartz, G.W., and Swygart, D. (2020). Circuits for Feature Selectivity in the Inner Retina. In The Senses: A Comprehensive Reference (Elsevier), pp. 275–292.

4. Seabrook, T.A., Burbridge, T.J., Crair, M.C., and Huberman, A.D. (2017). Architecture, Function, and Assembly of the Mouse Visual System. Annu. Rev. Neurosci. 40, 499–538.

5. Bos, R., Gainer, C., and Feller, M.B. (2016). Role for Visual Experience in the Development of Direction-Selective Circuits. Curr. Biol. 26, 1367–1375.

6. Wyatt, H.J., and Daw, N.W. (1975). Directionally sensitive ganglion cells in the rabbit retina: specificity for stimulus direction, size, and speed. J. Neurophysiol. 38, 613–626.

7. Yonehara, K., Shintani, T., Suzuki, R., Sakuta, H., Takeuchi, Y., Nakamura-Yonehara, K., and Noda, M. (2008). Expression of SPIG1 reveals development of a retinal ganglion cell subtype projecting to the medial terminal nucleus in the mouse. PLoS One 3.

8. Dhande, O.S., and Huberman, A.D. (2014). Retinal ganglion cell maps in the brain: Implications for visual processing. Curr. Opin. Neurobiol. 24, 133–142.

9. Sivyer, B., Tomlinson, A., and Taylor, W.R. (2019). Simulated Saccadic Stimuli Suppress ON-Type Direction-Selective Retinal Ganglion Cells via Glycinergic Inhibition. J. Neurosci. 39, 4312–4322.

10. Barlow, H.B., Hill, R.M., and Levick, W.R. (1964). Retinal ganglion cells responding selectively to direction and speed of image motion in the rabbit. J. Physiol. 173, 377–407.

11. Grzywacz, N.M., and Amthor, F.R. (2007). Robust directional computation in on-off directionally selective ganglion cells of rabbit retina. Vis. Neurosci. 24, 647–661.

12. Huberman, A.D., Wei, W., Elstrott, J., Stafford, B.K., Feller, M.B., and Barres, B.A. (2009). Genetic Identification of an On-Off Direction-Selective Retinal Ganglion Cell Subtype Reveals a Layer-Specific Subcortical Map of Posterior Motion. Neuron 62, 327–334.

13. Sivyer, B., Van Wyk, M., Vaney, D.I., and Taylor, W.R. (2010). Synaptic inputs and timing underlying the velocity tuning of direction-selective ganglion cells in rabbit retina. J. Physiol. 588, 3243–3253.

14. Lipin, M.Y., Taylor, W.R., and Smith, R.G. (2015). Inhibitory input to the direction-selective ganglion cell is saturated at low contrast. J. Neurophysiol. 114, 927–941.

15. Hoggarth, A., McLaughlin, A.J., Ronellenfitch, K., Trenholm, S., Vasandani, R., Sethuramanujam, S., Schwab, D., Briggman, K.L., and Awatramani, G.B. (2015). Specific wiring of distinct amacrine cells in the directionally selective retinal circuit permits independent coding of direction and size. Neuron 86, 276–291.

16. Grimes, W.N., Schwartz, G.W., and Rieke, F. (2014). The synaptic and circuit mechanisms underlying a change in spatial encoding in the retina. Neuron 82, 460–473.

17. Jacoby, J., Zhu, Y., DeVries, S.H., and Schwartz, G.W. (2015). An Amacrine Cell Circuit for Signaling Steady Illumination in the Retina. Cell Rep. 13, 2663–2670.

18. Mauss, A.S., Vlasits, A., Borst, A., and Feller, M. (2017). Visual Circuits for Direction Selectivity. Annu. Rev. Neurosci. 40, 211–230.

19. Frye, M. (2015). Elementary motion detectors. Curr. Biol. 25, R215–R217.

20. Borst, A., and Euler, T. (2011). Seeing Things in Motion: Models, Circuits, and Mechanisms. Neuron 71, 974–994.

21. Cooler, S., and Schwartz, G.W. (2021). An offset ON-OFF receptive field is created by gap junctions between distinct types of retinal ganglion cells. Nat. Neurosci. 24, 105–115.

22. Trenholm, S., Johnson, K., Li, X., Smith, R.G., and Awatramani, G.B. (2011). Parallel Mechanisms Encode Direction in the Retina. Neuron 71, 683–694.

23. Priebe, N.J. (2006). Tuning for Spatiotemporal Frequency and Speed in Directionally Selective Neurons of Macaque Striate Cortex. J. Neurosci. 26, 2941–2950.

24. Vaney, D.I., Sivyer, B., and Taylor, W.R. (2012). Direction selectivity in the retina: symmetry and asymmetry in structure and function. Nat. Rev. Neurosci. 13, 194–208.

25. Wei, W. (2018). Neural Mechanisms of Motion Processing in the Mammalian Retina. Annu. Rev. Vis. Sci. 4, 165–192.

26. Ding, H., Smith, R.G., Poleg-Polsky, A., Diamond, J.S., and Briggman, K.L. (2016). Species-specific wiring for direction selectivity in the mammalian retina. Nature 535, 105–110.

27. Matsumoto, A., Briggman, K.L., and Yonehara, K. (2019). Spatiotemporally Asymmetric Excitation Supports Mammalian Retinal Motion Sensitivity. Curr. Biol. 29, 3277–3288.e5.

28. Fried, S.I., Münch, T.A., and Werblin, F.S. (2002). Mechanisms and circuitry underlying directional selectivity in the retina. Nature 420, 411–414.

29. Taylor, W.R., and Vaney, D.I. (2002). Diverse Synaptic Mechanisms Generate Direction Selectivity in the Rabbit Retina. J. Neurosci. 22, 7712–7720.

30. Fried, S.I., Münch, T.A., and Werblin, F.S. (2005). Directional selectivity is formed at multiple levels by laterally offset inhibition in the rabbit retina. Neuron 46, 117–127.

31. Pei, Z., Chen, Q., Koren, D., Giammarinaro, B., Acaron Ledesma, H., and Wei, W. (2015). Conditional Knock-Out of Vesicular GABA Transporter Gene from Starburst Amacrine Cells Reveals the Contributions of Multiple Synaptic Mechanisms Underlying Direction Selectivity in the Retina. J. Neurosci. 35, 13219–13232.

32. Sivyer, B., and Williams, S.R. (2013). Direction selectivity is computed by active dendritic integration in retinal ganglion cells. Nat. Neurosci. 16, 1848–1856.

33. Jain, V., Murphy-Baum, B.L., Derosenroll, G., Sethuramanujam, S., Delsey, M., Delaney, K., and Awatramani, G.B. (2020). The functional organization of excitation and inhibition in the dendrites of mouse direction-selective ganglion cells. Elife 9.

34. Dhande, O.S., Estevez, M.E., Quattrochi, L.E., El-Danaf, R.N., Nguyen, P.L., Berson, D.M., and Huberman, A.D. (2013). Genetic Dissection of Retinal Inputs to Brainstem Nuclei Controlling Image Stabilization. J. Neurosci. 33, 17797–17813.

35. Rivlin-Etzion, M., Zhou, K., Wei, W., Elstrott, J., Nguyen, P.L., Barres, B.A., Huberman, A.D., and Feller, M.B. (2011). Transgenic Mice Reveal Unexpected Diversity of On-Off Direction-Selective Retinal Ganglion Cell Subtypes and Brain Structures Involved in Motion Processing. J. Neurosci. 31, 8760–8769.

36. Kretschmer, F., Tariq, M., Chatila, W., Wu, B., and Badea, T.C. (2017). Comparison of optomotor and optokinetic reflexes in mice. J. Neurophysiol. 118, 300–316.

37. Sakatani, T., and Isa, T. (2007). Quantitative analysis of spontaneous saccade-like rapid eye movements in C57BL/6 mice. Neurosci. Res. 58, 324–331.

38. Marques, T., Summers, M.T., Fioreze, G., Fridman, M., Dias, R.F., Feller, M.B., and Petreanu, L. (2018). A Role for Mouse Primary Visual Cortex in Motion Perception. Curr. Biol. 28, 1703–1713.e6.

39. Morrie, R.D., and Feller, M.B. (2018). A Dense Starburst Plexus Is Critical for Generating Direction Selectivity. Curr. Biol. 28, 1204–1212.e5.

40. Park, S.J.H., Borghuis, B.G., Rahmani, P., Zeng, Q., Kim, I.J., and Demb, J.B. (2015). Function and Circuitry of VIP+ Interneurons in the Mouse Retina. J. Neurosci. 35, 10685–10700.

41. Lee, S., Zhang, Y., Chen, M., and Zhou, Z.J. (2016). Segregated Glycine-Glutamate Co-transmission from vGluT3 Amacrine Cells to Contrast-Suppressed and Contrast-Enhanced Retinal Circuits. Neuron 90, 27–34.

42. Hanson, L., Sethuramanujam, S., DeRosenroll, G., Jain, V., and Awatramani, G.B. (2019). Retinal direction selectivity in the absence of asymmetric starburst amacrine cell responses. Elife 8, 1–20.

43. Oesch, N., Euler, T., and Taylor, W.R. (2005). Direction-selective dendritic action potentials in rabbit retina. Neuron 47, 739–750.

44. Emanuel, A.J., Kapur, K., and Do, M.T.H. (2017). Biophysical Variation within the M1 Type of Ganglion Cell Photoreceptor. Cell Rep. 21, 1048–1062.

45. Wienbar, S., and Schwartz, G.W. (2021). Differences in spike generation instead of synaptic inputs determine the feature selectivity of two retinal cell types. bioRxiv, 2021.10.19.464988.

46. Wehr, M., and Zador, A.M. (2003). Balanced inhibition underlies tuning and sharpens spike timing in auditory cortex. Nature 426, 860–863.

47. Schachter, M.J., Oesch, N., Smith, R.G., Taylor, W.R., and Rowland Taylor, W. (2010). Dendritic Spikes Amplify the Synaptic Signal to Enhance Detection of Motion in a Simulation of the Direction-Selective Ganglion Cell. PLoS Comput. Biol. 6, 1000899.

48. Jain, V., Hanson, L., Sethuramanujam, S., Gregg, R.G., Zhang, C., Smith, R.G., Berson, D., Mccall, M.A., Awatramani, G.B., and Awatramani, G. (2021). Gain control by sparse, ultra-slow glycinergic synapses. bioRxiv, 2021.05.03.442480.

49. Chen, Q., Pei, Z., Koren, D., and Wei, W. (2016). Stimulus-dependent recruitment of lateral inhibition underlies retinal direction selectivity. Elife 5, 1–19.

50. Kim, J.S., Greene, M.J., Zlateski, A., Lee, K., Richardson, M., Turaga, S.C., Purcaro, M., Balkam, M., Robinson, A., Behabadi, B.F., et al. (2014). Space–time wiring specificity supports direction selectivity in the retina. Nature 509, 331–336.

51. Lee, S., Kim, K., and Zhou, Z.J. (2010). Role of ACh-GABA Cotransmission in Detecting Image Motion and Motion Direction. Neuron 68, 1159–1172.

52. Matsumoto, A., Agbariah, W., Nolte, S.S., Andrawos, R., Levi, H., Sabbah, S., and Yonehara, K. (2021). Direction selectivity in retinal bipolar cell axon terminals. Neuron 109, 2928.

53. Hellmer, C.B., Hall, L.M., Bohl, J.M., Sharpe, Z.J., Smith, R.G., and Ichinose, T. (2021). Cholinergic feedback to bipolar cells contributes to motion detection in the mouse retina. Cell Rep. 37, 110106.

54. Strauss, S., Korympidou, M.M., Ran, Y., Franke, K., Schubert, T., Baden, T., Berens, P., Euler, T., and Vlasits, A.L. (2021). Center-surround interactions underlie bipolar cell motion sensing in the mouse retina. bioRxiv, 2021.05.31.446404.

55. Sethuramanujam, S., Matsumoto, A., deRosenroll, G., Murphy-Baum, B., McIntosh, J.M., Jing, M., Li, Y., Berson, D., Yonehara, K., and Awatramani, G.B. (2021). Rapid multi-directed cholinergic transmission in the central nervous system. Nat. Commun. 2021 121 12, 1–13.

56. Chen, Q., and Wei, W. (2018). Stimulus-dependent engagement of neural mechanisms for reliable motion detection in the mouse retina. J. Neurophysiol. 120, 1153–1161.

57. Zhang, C., and McCall, M.A. (2012). Receptor targets of amacrine cells. Vis. Neurosci. 29, 11–29.

58. Kim, T., Soto, F., and Kerschensteiner, D. (2015). An excitatory amacrine cell detects object motion and provides feature-selective input to ganglion cells in the mouse retina. Elife 4.

59. Lee, S., Chen, L., Chen, M., Ye, M., Seal, R.P., and Zhou, J. (2014). An unconventional glutamatergic circuit in the retina formed by vGluT3 amacrine cells. Neuron 84, 708–715.

60. Mani, A., Yang, X., Zhao, T., and Berson, D.M. (2021). A retinal circuit that vetoes optokinetic responses to fast visual motion. bioRxiv, 2021.10.31.466688.

61. Lien, A.D., and Scanziani, M. (2018). Cortical direction selectivity emerges at convergence of thalamic synapses. Nature 558, 80–86.

62. Wei, W., Hamby, A.M., Zhou, K., and Feller, M.B. (2011). Development of asymmetric inhibition underlying direction selectivity in the retina. Nature 469, 402–406.

63. Desai, N.S., Gray, R., and Johnston, D. (2017). A dynamic clamp on every rig. eNeuro 4, 1–17.

